# Odor imagery but not perception drives risk for food cue reactivity and increased adiposity

**DOI:** 10.1101/2023.02.06.527292

**Authors:** Emily E. Perszyk, Xue S. Davis, Jelena Djordjevic, Marilyn Jones-Gotman, Jessica Trinh, Zach Hutelin, Maria G. Veldhuizen, Leonie Koban, Tor D. Wager, Hedy Kober, Dana M. Small

## Abstract

Mental imagery has been proposed to play a critical role in the amplification of cravings. Here we tested whether olfactory imagery drives food cue reactivity strength to promote adiposity in 45 healthy individuals. We measured odor perception, odor imagery ability, and food cue reactivity using self-report, perceptual testing, and neuroimaging. Adiposity was assessed at baseline and one year later. Brain responses to real and imagined odors were analyzed with univariate and multivariate decoding methods to identify pattern-based olfactory codes. We found that the accuracy of decoding imagined, but not real, odor quality correlated with a perceptual measure of odor imagery ability and with greater adiposity changes. This latter relationship was mediated by cue-potentiated craving and intake. Collectively, these findings establish odor imagery ability as a risk factor for weight gain and more specifically as a mechanism by which exposure to food cues promotes craving and overeating.

## INTRODUCTION

The 21^st^ century rise in obesity coincides with the increased prevalence of palatable, energy-dense foods and ubiquitous cues signaling their availability^1^. Conditioned responses to food cues, such as increased salivation and brain responses, provide a measure of food cue reactivity. According to the ‘cued overeating model,’ such physiological and neural changes may be consciously experienced as craving, an intense desire for a particular food^2, 3^. Food cue reactivity is positively associated with body mass index (BMI)^4^ and highly predictive of weight change^5^.

One prominent theory of craving posits that repeated mental imagery of the sensory properties of a desired substance (e.g., food) leads to the intensification of cravings^6^.

Specifically, the Elaborated Intrusion Theory of Desire argues that craving episodes persist in a vicious cycle by which mental images provide immediate pleasure but exacerbate the awareness of a deficit and promote further planning to satisfy the desire^6^. Sensory imagery is a primary component of subjective food, drug, and alcohol craving, and the self-reported vividness of this mental imagery is positively associated with craving strength^7–13^. Accordingly, protocols in which individuals are asked to imagine palatable foods are frequently used to induce craving^14^.

Of central relevance to the current investigation, not all sensory modalities are similarly imaginable. The self-reported ability to imagine sights and sounds is nearly universal, whereas the ability to imagine odors and flavors varies widely across the population^15–19^. Previous work from our lab demonstrated that the self-reported vividness of imagined olfactory, but not visual, stimuli positively correlates with BMI^20^. These data raise the possibility that odor imagery ability confers risk for food cue reactivity and weight gain; however, whether this self-report measure reflects actual odor imagery ability is not clear. Also unknown is whether perceptual or neural measures of odor imagery ability are related to food cue reactivity, BMI, and weight gain susceptibility.

Odor imagery ability has been quantified as the extent to which imagining an odor decreases the detectability of a weak incongruent odor^21^. This ‘interference effect’ correlates with self-reported odor imagery ability in women^21^. It has also been used to identify good odor imagers (i.e., people with strong interference effects) who exhibit odor-imagery evoked increases in regional cerebral blood flow measured by positron emission tomography in primary and secondary olfactory regions^22^. However, since these regions are functionally heterogenous^23^, the correlation might reflect general processes like attention, saliency, and pleasantness, or more specific processes like odor quality coding. This distinction is important because imagery is based on the ability to reactivate sensory circuits that code the identity of the imagined stimulus^24^. In the case of olfaction, odor quality is encoded in patterns of activity across piriform cortex neurons^25–27^. In humans, these patterns can be decoded with multi-voxel pattern analyses (MVPA) of functional magnetic resonance imaging (fMRI) data^28^. Whether imagining an odor reactivates these odor quality patterns has not been tested.

In the current study, we set out to first determine if the interference effect – a performance-based perceptual measure of odor imagery ability – is associated with self-reported ability and the decoding of odor quality from fMRI patterns evoked by real and/or imagined odors in the piriform cortex (Fig. 1a). Our second goal was to test if the perceptual (i.e., the interference effect) and neural (i.e., piriform decoding of imagined odors) measures of odor imagery ability are associated with behavioral food cue reactivity, quantified as cue-induced craving and cue-potentiated food intake (Fig. 1b). We also explored whether imagining odors elicits an independently established brain measure of food cue reactivity (Fig. 1b), the Neurobiological Craving Signature (NCS). The NCS is a recently developed multivariate brain activity pattern, or neuromarker, that predicts the intensity of self-reported food and drug craving across distinct samples^29^. Finally, we sought to test if odor imagery is associated with current or change in adiposity over one year (Fig. 1b). We hypothesized that better odor imagery ability would be associated with stronger food cue reactivity and greater change in adiposity (Fig. 1c), with food cue reactivity mediating the association between odor imagery and adiposity change (Fig. 1d).

**Fig. 1:**
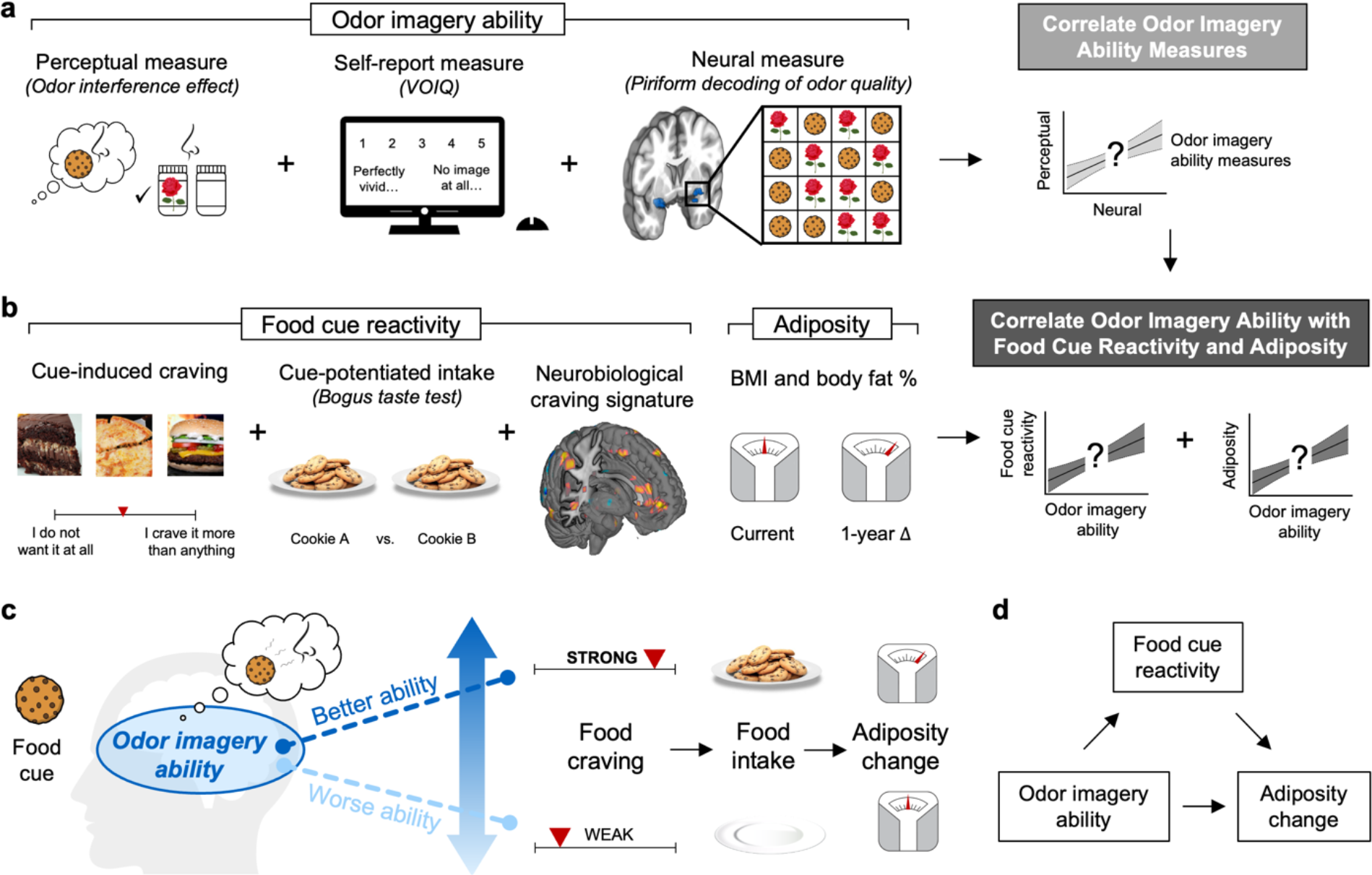
Study Overview and Model. (a) Our first goal was to establish relationships between three measures of odor imagery ability: a validated perceptual measure adapted from Djordjevic et al. (2004)^21^, a self-report measure (the Vividness of Olfactory Imagery Questionnaire or VOIQ^30^), and a new neural measure based upon the piriform decoding of odor quality. See Figs. 2–4 for additional details on the perceptual and neural measures. (b) Our second goal was to correlate odor imagery ability with two behavioral measures of food cue reactivity: cue-induced craving from an established paradigm^31^ and cue-potentiated intake in a bogus taste test^32^. We also examined the extent to which smelling versus imagining odors elicits an independently established brain measure of food cue reactivity, the Neurobiological Craving Signature^29^. Our third goal was to test the associations between odor imagery ability and both current and one-year changes in adiposity. (c) We hypothesized that in response to learned food cues, individuals with a better ability to imagine odors would experience stronger cravings that compel them to overeat and gain weight. In contrast, individuals with a worse ability to imagine odors would experience weaker cravings that have a low impact on their eating and weight. (d) We predicted that food cue reactivity would mediate the association between odor imagery ability and adiposity change, such that odor imagery indirectly affects adiposity change via a food cue reactivity-dependent mechanism.

To test these hypotheses, we collected data from 45 adults (ages 18 – 42 years) with a range of BMIs (18.32 – 53.44 kg/m^2^). Participants completed three behavioral sessions and an fMRI scan at baseline to quantify odor imagery ability and food cue reactivity. They returned one year later for a follow-up session to assess adiposity change. As predicted, stronger interference effects were associated with better decoding accuracies of imagined, but not real, odors in the piriform cortex. Decoding also correlated positively with food intake. Most importantly, food craving and intake mediated the relationships between odor imagery ability and changes in BMI and body fat percentage, respectively. Collectively, these findings establish odor imagery ability as a risk factor for weight gain susceptibility and more specifically as a mechanism by which exposure to food cues promotes food craving and subsequent intake.

## RESULTS

### Self-Report and Perceptual Measures of Odor Imagery Ability are Correlated

To assess subjective experience of the ability to imagine odors and flavors, we used the Vividness of Olfactory Imagery Questionnaire (VOIQ)^30^ and the Vividness of Food Imagery Questionnaire (VFIQ)^20^, respectively. Our performance-based perceptual measure was adapted from Djordjevic et al. (2004)^21^ and is detailed in the Materials and Methods section (and see Fig. 2a). In brief, participants were instructed to imagine the smell or sight of a rose or cookie while trying to determine which of two samples contained either the same odor (matched trial) or the other odor (mismatched trial) at their detection threshold level (determined prior to the test). In the no imagery condition, odor detection trials were performed in the absence of imagery. The interference effect (i.e., perceptual measure of odor imagery ability) was calculated by subtracting detection accuracy (% trials correct) in mismatched trials from that in matched trials of the odor imagery condition.

**Fig. 2:**
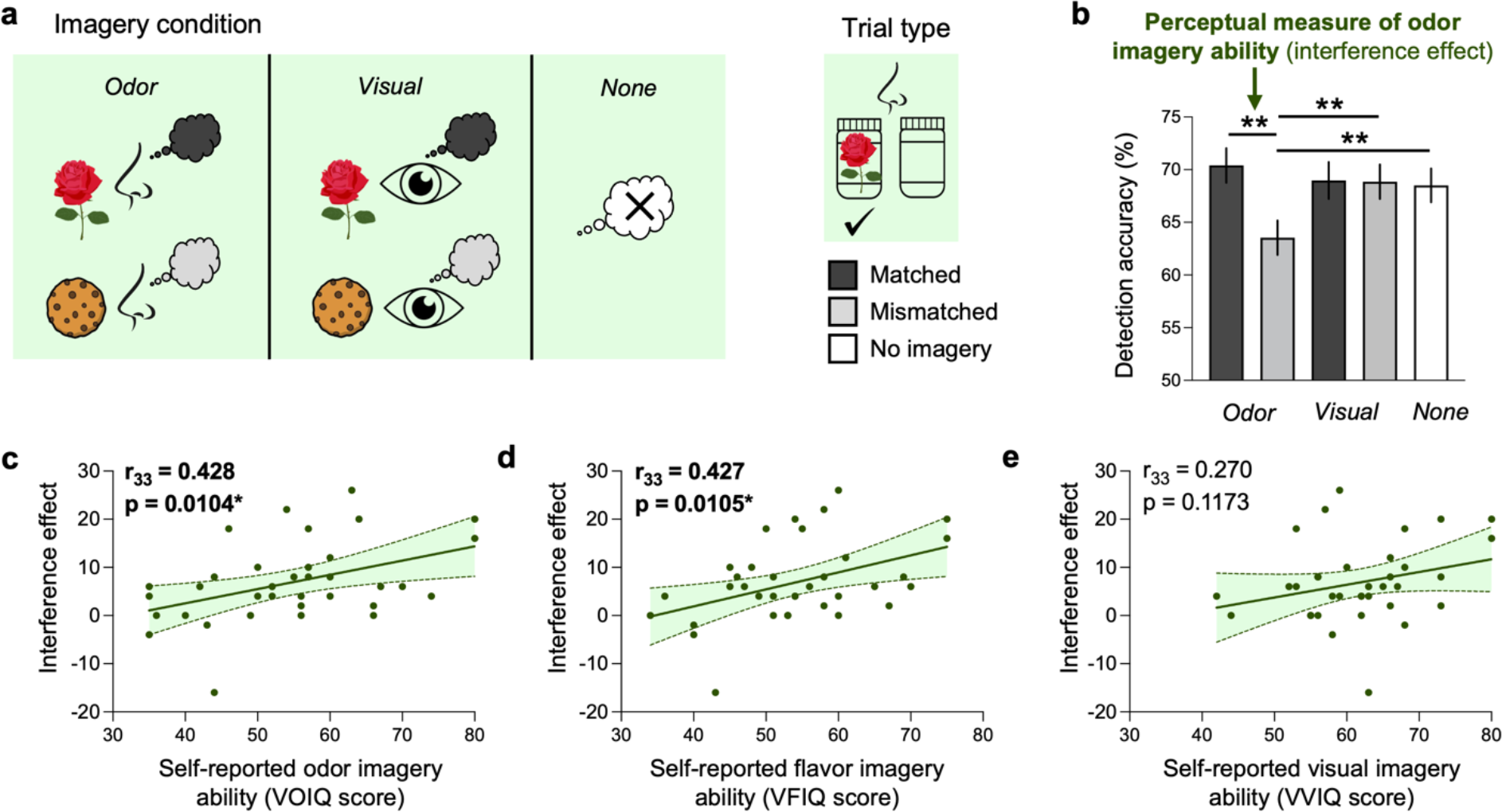
The Perceptual Measure of Odor Imagery Ability Correlates Positively with Self-Reported Odor and Flavor, but not Visual, Imagery Ability. (a) In the adapted perceptual task^21^ to quantify odor imagery ability, participants were instructed to imagine the smell or sight of a rose/cookie or nothing at all while trying to detect either the same (matched trial) or the other (mismatched trial) odor at their threshold level. (b) An ANOVA revealed a significant imagery condition × trial type interaction on detection accuracy. This effect was a result of the interference (rather than facilitation) of odor imagery on detection, such that performance on mismatched trials was significantly worse during odor imagery than in the visual or no imagery conditions. (c–e) The perceptual measure of odor imagery ability (i.e., the interference effect) positively correlated with self-reported odor (c) and flavor (d), but not visual (e), imagery ability. Bar plots represent M ± SEM. Fitted scatterplots depict single participants and the 95% CI. VOIQ, Vividness of Olfactory Imagery Questionnaire^30^; VFIQ, Vividness of Food Imagery Questionnaire^20^; VVIQ, Vividness of Visual Imagery Questionnaire^17^. *p < 0.05; **post-hoc comparison p < 0.0167 (0.05 / 3 tests).

As in previous work^21^, we found a significant interaction between imagery condition (odor/visual) and trial type (matched/mismatched) on detection accuracy after controlling for odor type (rose/cookie; F_1,275_ = 6.270, p = 0.0129). This was driven by worse performance on mismatched compared to matched trials during odor (t_137_ = 3.870, p = 0.0002), but not visual (t_137_ = 0.055, p = 0.9560) imagery (Fig. 2b). We next tested whether facilitation during matched trials, interference during mismatched trials, or a combination of the two contributed to this effect. We observed no impact of imagery condition on detection accuracy in matched versus no imagery trials (F_1,207_ = 2.926, p = 0.0886). In contrast, there was a main effect of imagery on detection across mismatched and no imagery trials (F_1,207_ = 6.187, p = 0.0137). Follow-up pairwise comparisons revealed that participants performed worse during odor mismatched versus visual mismatched trials (t_137_ = 2.712, p = 0.0076) and during odor mismatched versus no imagery trials (t_137_ = 2.434, p = 0.0162). There was no difference in visual mismatched versus no imagery trials (t_137_ = 0.163, p = 0.8709). Collectively, these data replicate prior findings showing that odor imagery impairs mismatched odor detection without improving matched detection^21^.

To determine whether this perceptual measure corresponded to self-reported odor imagery ability, we correlated the interference effect with perceived vividness of imagined odors (VOIQ^30^) and flavors (VFIQ^20^). Both correlations were significant (Fig. 2c and 2d). By contrast, no significant association was observed between the interference effect and self-reported visual imagery (Fig. 2e) in the Vividness of Visual Imagery Questionnaire (VVIQ)^17^. Similarly, the difference in detection accuracy on matched versus mismatched trials of the visual imagery condition did not correlate with self-reported odor (r_33_ = 0.306, p = 0.0735), flavor (r_33_ = 0.247, p = 0.1519), or visual (r_33_ = 0.155, p = 0.3742) imagery ability. The self-report and perceptual measures of odor imagery ability also did not vary by sex, age, household income, olfactory function, odor ratings, sniff parameters, hunger, or typical consumption of unhealthy foods (Supplementary Tables 1 and 2). These results confirm that the self-report and perceptual measures are associated, supporting the validity of using the interference effect as a measure of odor imagery ability.

### Imagining and Smelling Odors Activate Partly Overlapping Brain Regions

To assess brain responses to real odors, rose and cookie odors (or clean air) were repeatedly delivered via an olfactometer at moderate intensity to participants undergoing fMRI scanning. These trials were interspersed with ones in which participants were instructed to imagine the odors while sniffing clean air (Fig. 3a). As there was no main effect of odor type (rose/cookie) on fMRI activity (all p_FWE_ [family-wise error corrected] ≥ 0.3214), we collapsed across the odorants in the subsequent univariate analyses. Consistent with previous studies, we observed a main effect of smelling odors > smelling clean air in the bilateral insula, piriform/amygdala, orbitofrontal cortices, cerebellum, and middle frontal and cingulate gyri, along with the right thalamus and supramarginal gyrus and the left pre- and postcentral gyri (Fig. 3b, Supplementary Table 3). Many of the same regions were responsive to imagining odors > imagining clean air, including the bilateral insula, right putamen, and left cerebellum (Fig. 3c, Table 1). Given that most prior neuroimaging studies on odor imagery have contrasted imagining odors > smelling clean air, we also tested this effect. We found significant responses in the bilateral insula, putamen extending into the piriform cortices, pallidum, and orbitofrontal, middle frontal, and precentral gyri, along with the left cerebellum and the right hippocampus and postcentral, supramarginal, and cingulate gyri (Extended Data Fig. 1, Supplementary Table 4).

**Fig. 3:**
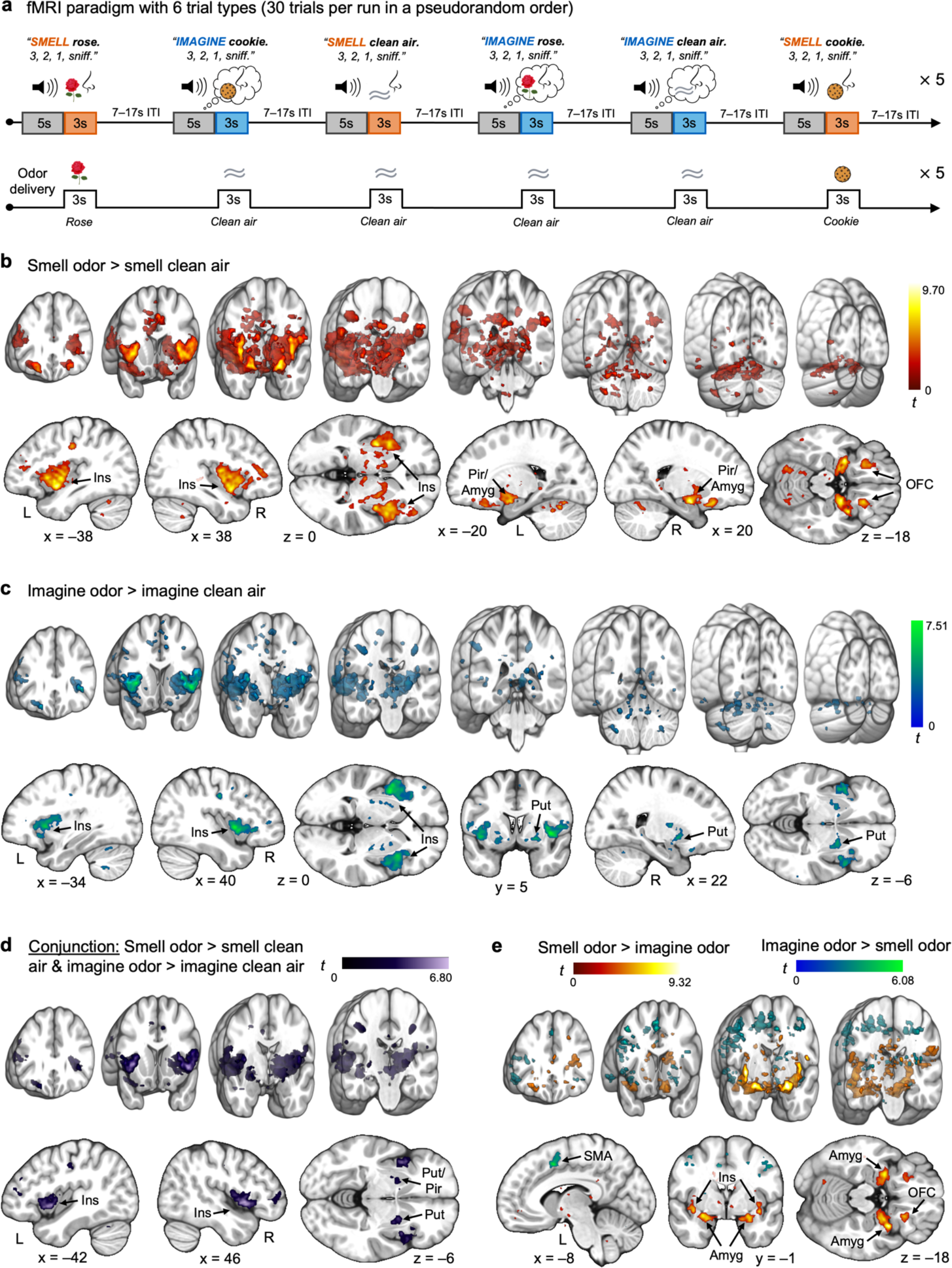
Univariate fMRI Activity During Odor Imagery Partly Mimics that During Real Odor Perception. (a) Overview of the fMRI paradigm with five scan runs. Five presentations each of six trial types (30 total) were pseudorandomized per run. Trials began with a 5s auditory cue including the trial type (e.g., “smell rose”) and a sniff countdown of “3, 2, 1, sniff.” In the “smell” trials, participants sniffed during the 3s delivery of a rose or cookie odor or clean air via an MRI-compatible olfactometer. During the “imagine” trials, they sniffed during a 3s clean air delivery. Trials were separated by an intertrial interval (ITI) of 7–17s (mean = 10s). (b) BOLD responses to smelling odors (rose and cookie) > smelling clean air were significant in the bilateral insula, piriform/amygdala, orbitofrontal cortices, cerebellum, and middle frontal and cingulate gyri, among other regions. (c) BOLD responses to imagining odors > imagining clean air (while sniffing) were significant in the bilateral insula, right putamen, and left cerebellum. (d) BOLD responses in the conjunction of smelling odors > smelling clean air and imagining odors > imagining clean air were significant in the bilateral insula and putamen extending into the piriform cortices, along with the left precentral gyrus. (e) BOLD responses to smelling odors > imagining odors were significant in the bilateral insula and amygdala and the right uncus and orbitofrontal cortex, among other regions. Those to imagining odors > smelling odors were significant in the left supplementary motor area. Brain sections show the SPM *t*-map (*p*uncorrected < 0.005, clusters of at least 5 voxels) overlaid onto an anatomical template in MNI coordinates for illustrative purposes. In each panel, the top row depicts 3D coronal sections (18mm thick) evenly spanning y = 56 to –88mm, and the bottom row highlights important areas of activation with custom coordinates (see Table 1 and Supplementary Table 3). Color bars depict *t* values. L, left; R, right; Amyg, amygdala; Ins, insula; OFC, orbitofrontal cortex; Pir, piriform cortex; Put, putamen.

**Table 1.**
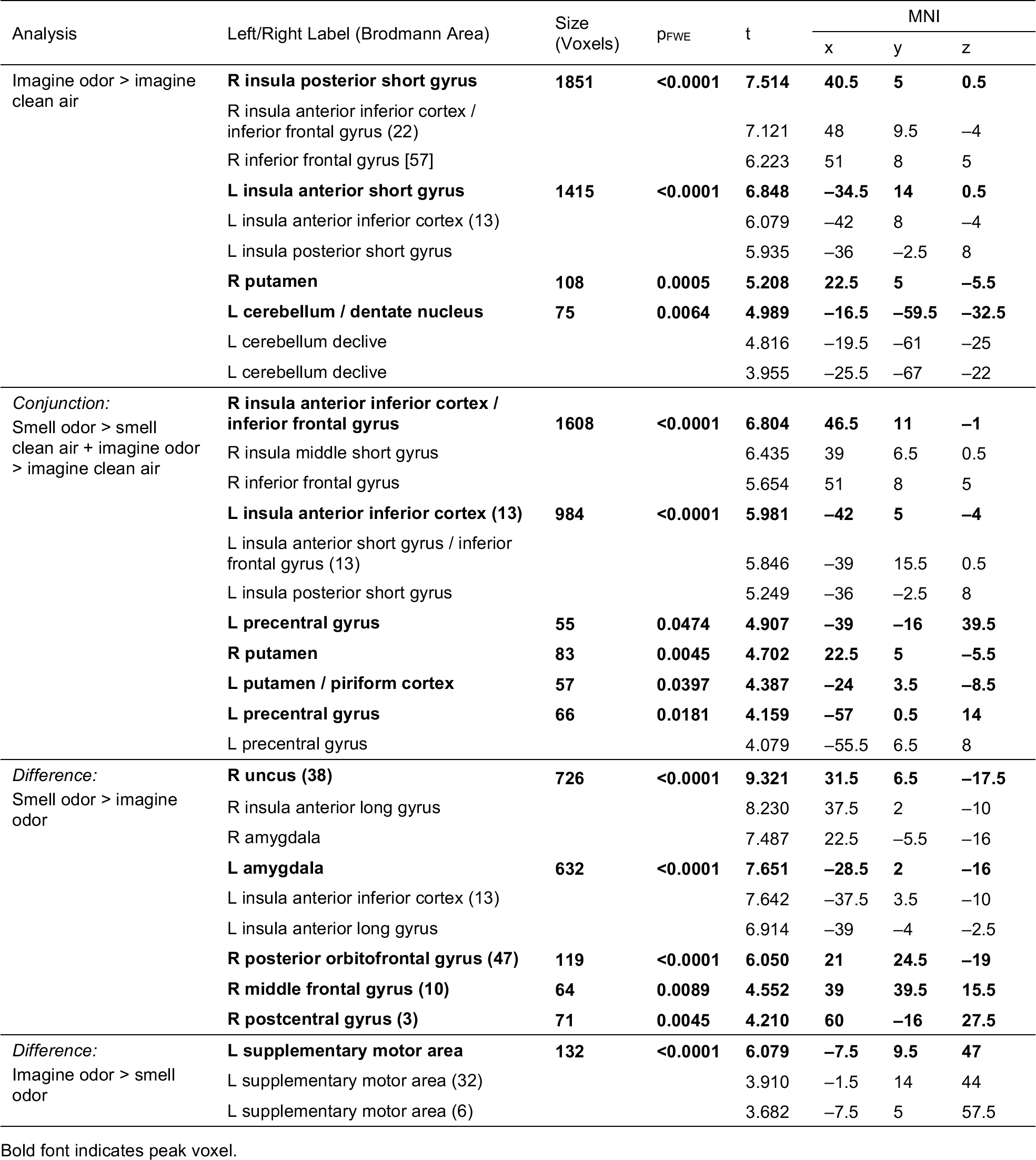
Brain Regions with Significant Responses During Odor Imagery Versus Perception.

To isolate areas of activation common to smelling and imagining odors, the comparisons of smelling odors > smelling clean air and imagining odors > imagining clean air were entered into a conjunction analysis using the conjunction null hypothesis. This revealed common activations in the bilateral insula and putamen extending into the piriform cortices, as well as in the left precentral gyrus (Fig. 3d, Table 1). We also compared the differences of smelling odors > imagining odors and of imagining odors > smelling odors to isolate clusters specific to perception versus imagery and vice versa. The bilateral insula and amygdala, right uncus, and right lateral orbitofrontal, middle frontal, and postcentral gyri were more responsive to real odors, whereas the left supplementary motor area (SMA) showed a stronger response for imagined odors (Fig. 3e, Table 1). These analyses confirmed that as in odor perception, odor imagery engages brain areas critical for olfactory processing, such as the piriform and insular cortices.

Lastly, we regressed the perceptual measure of odor imagery ability (i.e., the interference effect) against whole-brain univariate BOLD responses during odor imagery. We did not observe any effects for imagining odors > smelling clean air. By contrast, the perceptual measure of odor imagery ability was negatively associated with brain response to imagining odors > imagining clean air in the right fusiform gyrus (t_42_ = 5.038, p_FWE_ < 0.0001, size = 113 voxels, x = 28, y = –62, z = –14). We did not find any significant relationships in the piriform cortex, including after small-volume correction. These results suggest that odor imagery ability in the current study may not correspond to the magnitude of imagined odor-evoked activity in the primary olfactory cortex. They do not, however, indicate whether odor imagery ability is associated with odor quality coding in this region. This is important because odor quality is coded across distributed patterns of activation rather than reflected in average univariate responses^28^.

### Piriform Decoding of Imagined, but Not Actual, Odors Correlates with the Perceptual Measure of Odor Imagery Ability

To isolate fMRI patterns specific to odor quality coding, we performed MVPA in left and right piriform cortex regions of interest (ROIs; Fig. 4a). We first trained and tested a support vector machine (SVM) on the voxel-based patterns of activation evoked while smelling the odors using a leave-one-run-out, cross-validated approach per individual (Fig. 4b). This analysis revealed significant group-level decoding in the right piriform cortex (mean accuracy = 63.2%, chance = 50%, t_43_ = 2.991, p = 0.0046; Fig. 4c), along with greater decoding accuracies in the right compared to the left piriform cortices (t_43_ = 2.407, p = 0.0205). Next, we examined whether the imagined odor qualities also activated distributed neural patterns by training and testing the SVM on the voxel-based patterns of activation evoked during imagery of the two odor qualities. We did not observe significant group-level decoding in either ROI (Fig. 4c). Likewise, crossmodal decoding (training on real odors and testing on imagined odors, and vice versa) did not produce any significant effects (Fig. 4c).

**Fig. 4:**
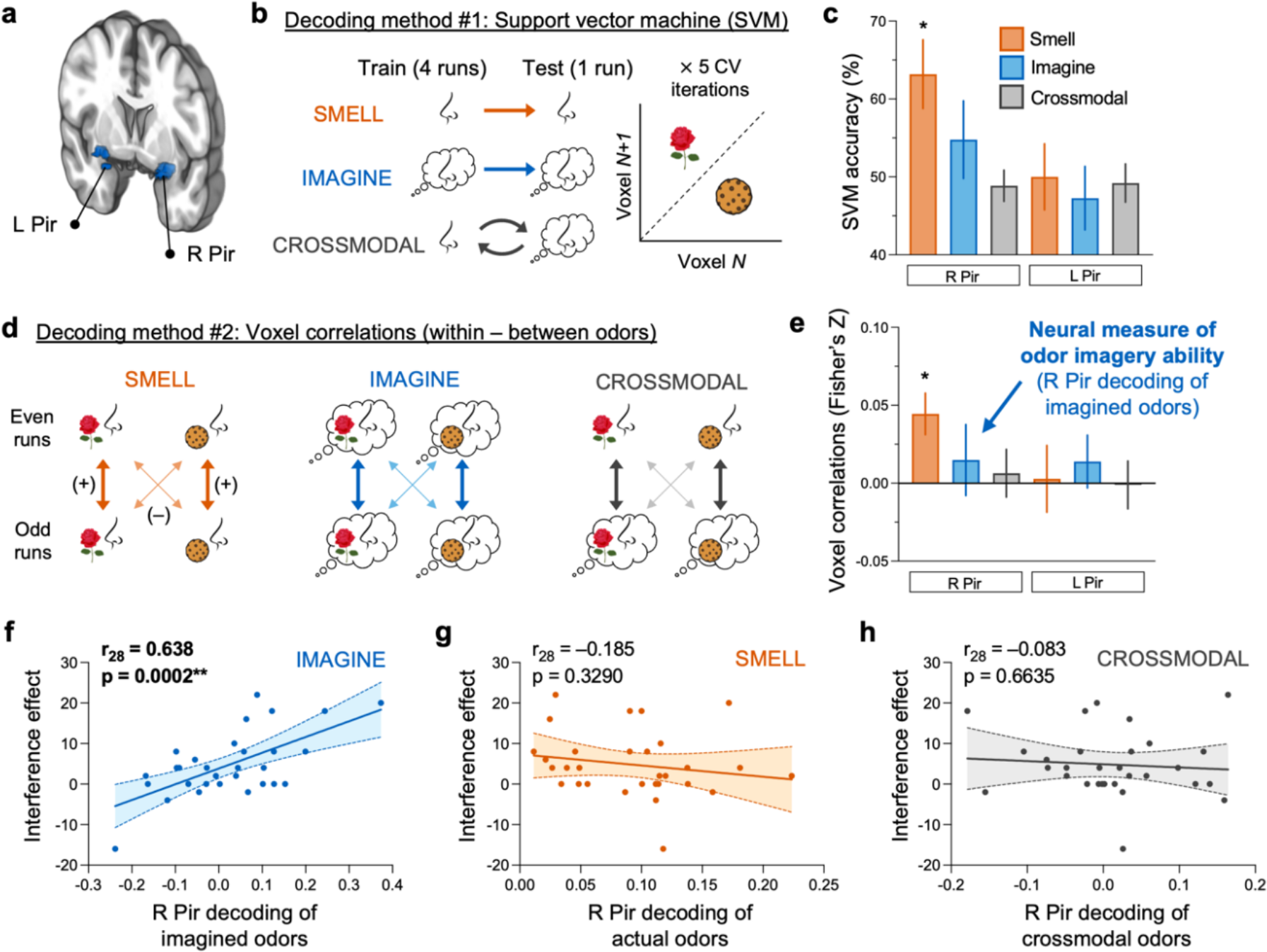
Decoding of Imagined, but Not Actual, Odors in the Right Piriform Cortex Provides a Neural Measure of Odor Imagery Ability. (a) Regions of interest for the neural decoding analyses. (b) SVMs were trained to classify rose versus cookie using data from four runs and tested on data from the fifth left-out run across five CV iterations. In the “smell odor” and “imagine odor” conditions, SVMs were trained and tested on voxel patterns from the same modality (actual odors and imagined odors, respectively). For crossmodal decoding, the SVM was trained on real odor patterns and tested on imagined odor patterns (and vice versa). (c) SVM accuracies for smelling actual odors in the right piriform cortex were significant at the group-level. (d) Split-half Fisher’s Z-transformed voxel correlations calculated between odor (e.g., smelling the rose odor in even runs versus smelling the cookie odor in odd runs) were subtracted from those calculated within odor (e.g., smelling the rose odor in even versus odd runs) as a more sensitive index of neural decoding. (e) Voxel correlations for smelling actual odors in the right piriform cortex were significant at the group-level. (f–h) The perceptual measure of odor imagery ability (i.e., the interference effect) positively correlated with right piriform decoding of imagined (f), but not real (g) or crossmodal (h), odors using voxel correlations (decoding method #2). Bar plots represent M ± SEM. Fitted scatterplots depict single participants and the 95% CI. L, left; R, right; Ins, insula, Pir, piriform; CV, cross-validation. *p < 0.01; **p < 0.001.

Leave-one-run-out cross-validation in which an SVM is trained and tested on the average run-wise parameter estimates for each condition provides a relatively insensitive outcome metric. For five scan runs, one decoding error reduces the accuracy estimate by 20%, such that the read-out for any given participant is a multiple of this number (i.e., either 20, 40, 60, 80, or 100%). We therefore employed a more sensitive decoding method by analyzing the split-half voxel correlations for the within-odor (e.g., smelling rose in even runs versus smelling rose in odd runs) minus the between-odor (e.g., smelling rose in even runs versus smelling cookie in odd runs) voxel-based activity patterns (Fig. 4d). In line with our SVM analyses, we performed separate voxel correlations for real, imagined, and crossmodal odors. Again, decoding accuracy was only significant for smelling real odors in the right piriform cortex (t_43_ = 3.342, p = 0.0017; Fig. 4e). Given that odor imagery ability varies widely across the population_18_, the lack of main effect of imagined odor decoding is unsurprising. However, the re- activation of sensory codes during imagery may occur in those individuals with vivid imagery. In this case, decoding of imagined odor qualities in the piriform cortex should correlate with the self-reported and perceptual measures of odor imagery ability.

To test this, we next examined the relationships between olfactory decoding (using the split-half voxel correlations method) and the interference effect. We restricted our analyses to the right piriform cortex in the 30 individuals with discriminable neural patterns for actual odors to ensure that any effects would not be driven by an inability to decode altogether. We observed a strong positive association between right piriform decoding of imagined odors and our perceptual measure of odor imagery ability (i.e., the interference effect; Fig. 4f). Similar analyses using the self-report measures of odor (p = 0.0571) and flavor (p = 0.0722) imagery ability approached significance. In contrast, there were no significant associations between the perceptual or self-report measures of odor imagery ability and the fMRI patterns evoked during actual odor presentations (Fig. 4g) or in the crossmodal datasets (Fig. 4h). The results remained largely unchanged when including the full sample (n = 44; Supplementary Table 5). Imagined odor decoding was also unrelated to the demographic variables, olfactory function, odor ratings, sniff parameters, hunger, and typical consumption of unhealthy foods (Supplementary Table 1). Collectively, these data demonstrate that odor imagery ability is associated with successful activation of distinct imagined odor quality codes in the right piriform cortex.

### Odor Imagery Ability is Associated with Stronger Food Cravings for Liked Foods

To test our prediction that odor imagery ability is associated with food cue reactivity, we used a measure of cue-induced food craving in which participants were asked to rate the strength of their craving in response to the presentation of 90 palatable food images^31^ (see example stimuli in Fig. 1b). We found no significant relationships between the perceptual measure of odor imagery ability (i.e., the interference effect; Fig. 5a) or the neural measure of odor imagery ability (i.e., right piriform decoding of imagined odors; Fig. 5b) and the average rating of food craving strength. Likewise, the decoding of actual odors in the right piriform cortex was unrelated to food craving (Fig. 5c).

**Fig. 5:**
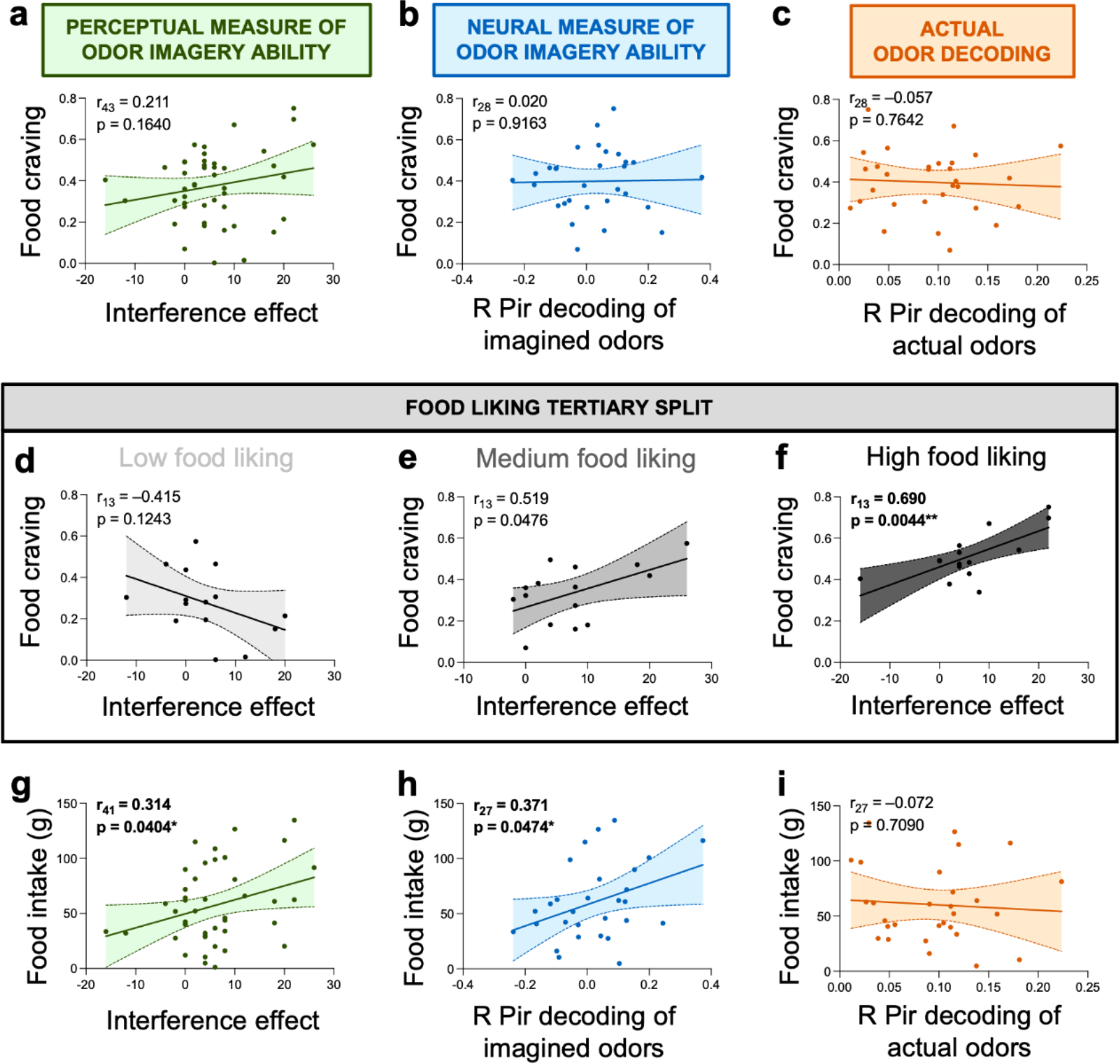
Odor Imagery Ability Contributes to Food Cue Reactivity. (a–c) Food craving did not correlate with the perceptual (a) or neural (b) measures of odor imagery ability or with actual odor decoding (c). (d–f) There was a significant interaction between food liking and the perceptual measure of odor imagery ability on craving (p = 0.0057). Following a tertiary split to separate participants by their average food liking, the interference effect was unrelated to food craving in the low (d) and medium (e) food liking groups after Bonferroni correction for the three tests performed. By contrast, there was a positive correlation in the high food liking group (f). (g–i) Food intake positively correlated with the perceptual (g) and neural (h) measures of odor imagery ability, but not with actual odor decoding (i). Fitted scatterplots depict single participants and the 95% CI. R, right; Pir, piriform; LHS, Labeled Hedonic Scale^33^. *p < 0.05; **p < 0.01.

However, the rated liking of the foods depicted in the pictures was variable and significantly correlated with craving (Supplementary Table 6). We therefore reasoned that odor imagery may intensify cravings specifically for foods that are liked and constructed a linear regression model to test for the presence of an interaction between odor imagery and food liking on the average craving rating. As predicted, the interaction was significant for the perceptual measure of odor imagery ability (t_41_ = 2.918, p = 0.0057) and approached significance for the neural measure (t_26_ = 1.835, p = 0.0780). For graphical purposes and to better understand the nature of this interaction, we used a tertiary split (n = 15 each) to separate participants based on their average food liking rated on the Labeled Hedonic Scale (LHS)^33^. In the low liking group (mean LHS rating = –0.17, range = –66.60 to 11.68), there was no correlation between the perceptual measure of odor imagery ability and food craving (Fig. 5d). In the medium liking group (mean LHS rating = 19.52, range = 11.83 to 27.98), a positive trend emerged that was not significant after correction for multiple comparisons (Fig. 5e). In contrast, the high liking group (mean LHS rating = 37.69, range = 29.85 to 48.98) showed a strong positive association even after correction for multiple comparisons (Fig. 5f).

We also performed a follow-up analysis using a linear mixed effects model with the individual ratings for each of the 90 food pictures rather than participant averages. Craving was designated as the outcome variable; the interference effect, food liking, and the interaction of the two as fixed effects; and participant as a random effect. Testing this model once again revealed a significant interaction effect (F_1,3996_ = 7.571, p = 0.0060) whereby cravings for liked but not disliked foods were more intense in individuals with vivid odor imagery. In addition, accounting for subjective hunger ratings – which were positively correlated with food craving (Supplementary Table 6) – did not impact any of the results. Collectively, these data suggest that good odor imagery ability paired with high food liking may give rise to intense food cravings.

### Odor Imagery Ability Correlates with Cue-Potentiated Food Intake

Having identified a role for odor imagery ability in food craving, we next tested for associations with our second measure of food cue reactivity, cue-potentiated food intake. We performed a validated bogus taste test^32^ in which participants were presented with two plates of cookies. They were instructed to sample as much as they wanted while responding to questions about the sensory properties of the cookies. They were not told that the real purpose of the test was to quantify how much was consumed in grams. In line with our expectations, the perceptual (r_41_ = 0.314, p = 0.0404; Fig. 5g) and neural (r_27_ = 0.371, p = 0.0474; Fig. 5h) measures of odor imagery ability were each positively associated with food intake. Additionally, we performed separate linear regressions to adjust for sex – since males ate more than females – and cookie liking ratings, which were positively correlated with the amount consumed (Supplementary Table 6). Both the perceptual (β = 0.351, p = 0.0098) and neural (β = 0.420, p = 0.0136) measures of odor imagery ability remained as significant predictors of intake in these models. Interestingly, right piriform decoding of actual odors was unrelated to cookie consumption (r_27_ = –0.072, p = 0.7090; Fig. 5h). These findings indicate that the association is specific to odor quality codes evoked during imagery.

### Neurobiological Craving Signature Responses are Stronger while Imagining a Food Versus Nonfood Odor

To build upon our use of two behavioral food cue reactivity measures, our next step was to explore the extent to which smelling versus imagining odors elicits brain food cue reactivity. We assessed brain food cue reactivity using the Neurobiological Craving Signature (NCS), a multivariate brain pattern that reliably predicts self-reported drug and food craving across independent samples^29^. We tested a version of the NCS trained only on visual food cues and identified the ‘pattern response’ value for each participant and fMRI contrast in the current study. This pattern response describes the similarity of the participant’s contrast image (e.g., while imagining odors) to the NCS and therefore the predicted level of food craving for that individual. As such, greater NCS pattern responses indicate stronger similarity to the craving map and higher predicted levels of food craving. Although the NCS is capable of distinguishing drug users from non-users in prior work, it was not primarily trained to detect individual differences in craving. NCS pattern responses also partially depend on factors such as overall fMRI signal^34^, which differed in coverage of the parietal lobe across participants in the current study (Extended Data Fig. 2a). However, NCS pattern responses are particularly well suited for assessing the impacts of within-subject interventions or contrasting contexts (e.g., examining their modulation by condition, such as smelling versus imagining the odor types and clean air).

In testing the latter, we found significant main effects of perceptual modality (smelling/imagining; F_1,258_ = 7.765, p = 0.0057) and odor type (rose/cookie/clean air; F_2,258_ = 9.716, p < 0.0001) and an interaction of the two (F_2,258_ = 4.100, p = 0.0177; Extended Data Fig. 2b) on NCS pattern responses. Follow-up comparisons revealed significantly greater NCS pattern responses to smelling versus imagining the cookie (t_86_ = 3.192, p = 0.0020) and rose (t_86_ = 4.593, p < 0.0001) odors, with no difference in smelling versus imagining clean air (t_86_ = 0.376, p = 0.7077). Smelling both the rose (t_86_ = 3.078, p = 0.0028) and cookie (t_86_ = 3.862, p = 0.0002) odors resulted in greater NCS pattern responses than smelling clean air. In contrast, neither imagining the cookie (t_86_ = 0.753, p = 0.0832) nor the rose (t_86_ = 0.499, p = 0.6193) odor yielded greater NCS pattern responses than imagining clean air. However, imagining the cookie odor did elicit stronger NCS pattern responses than imagining the rose odor (t_86_ = 3.068, p = 0.0029), an effect that did not occur for smelling the cookie versus rose odor (t_86_ = 0.428, p = 0.6695).

These discrepancies suggest that the brain signature for craving is weaker during odor imagery than during real perception. Yet it may also be more finely tuned to food versus nonfood cues, such that the craving level predicted by the NCS is greater for food odors than nonfood odors during olfactory imagery.

### Odor Imagery Ability is Not Related to Current Adiposity

In contrast with prior work^20^, the self-reported, perceptual, and neural measures of odor imagery ability were not significantly associated with current adiposity defined by BMI or body fat percentage (Supplementary Table 1). However, we speculated that the variance in BMI within the current participant sample (BMI: M = 26.12, SD = 6.81, Range = 18.32–53.44 kg/m^2^) may have differed from that across the two experiments (BMI: M = 25.75, SD = 5.06, Range = 17.70–38.70 kg/m^2^) in the previous study comparing VOIQ score with BMI^20^. To test this, we used a two-sample F-test for equal variances and found that the BMI variance in the present study was significantly greater than in the prior study (F44,81 = 1.810, p = 0.0210). As the causes of obesity are heterogeneous, one possible explanation for the lack of a significant correlation between odor imagery ability and current adiposity here could be the inclusion of participants across a wider range of BMIs.

### Food Cue Reactivity Mediates the Relationship between Odor Imagery Ability and Adiposity Change

In the previous analyses, we demonstrate that odor imagery ability is associated with food cue reactivity but not current adiposity. To test our overarching hypothesis that odor imagery ability intensifies food cravings and increases consumption to promote longer-term weight gain, we first used correlation analyses to assess the relationships among these variables. Neither measure of odor imagery ability was significantly correlated with changes in BMI or body fat percentage over one year from the baseline to follow-up sessions (Fig. 6a-d). Food craving in the cue-induced craving paradigm was also not associated with cue-potentiated food intake in the bogus taste test (r_41_ = 0.255, p = 0.0983). However, food craving predicted changes in BMI (Fig. 6e) but not body fat percentage (r_41_ = 0.229, p = 0.1402), whereas food intake predicted changes in body fat percentage (Fig. 6f) but not BMI (r_39_ = 0.263, p = 0.0964). Accounting for age – which was positively correlated with change in BMI (Supplementary Table 7) – did not impact any of the results. Additionally, changes in adiposity were unrelated to sex, household income, olfactory function, food liking, typical consumption of unhealthy foods, and changes in physical activity over the year (Supplementary Table 7).

**Fig. 6:**
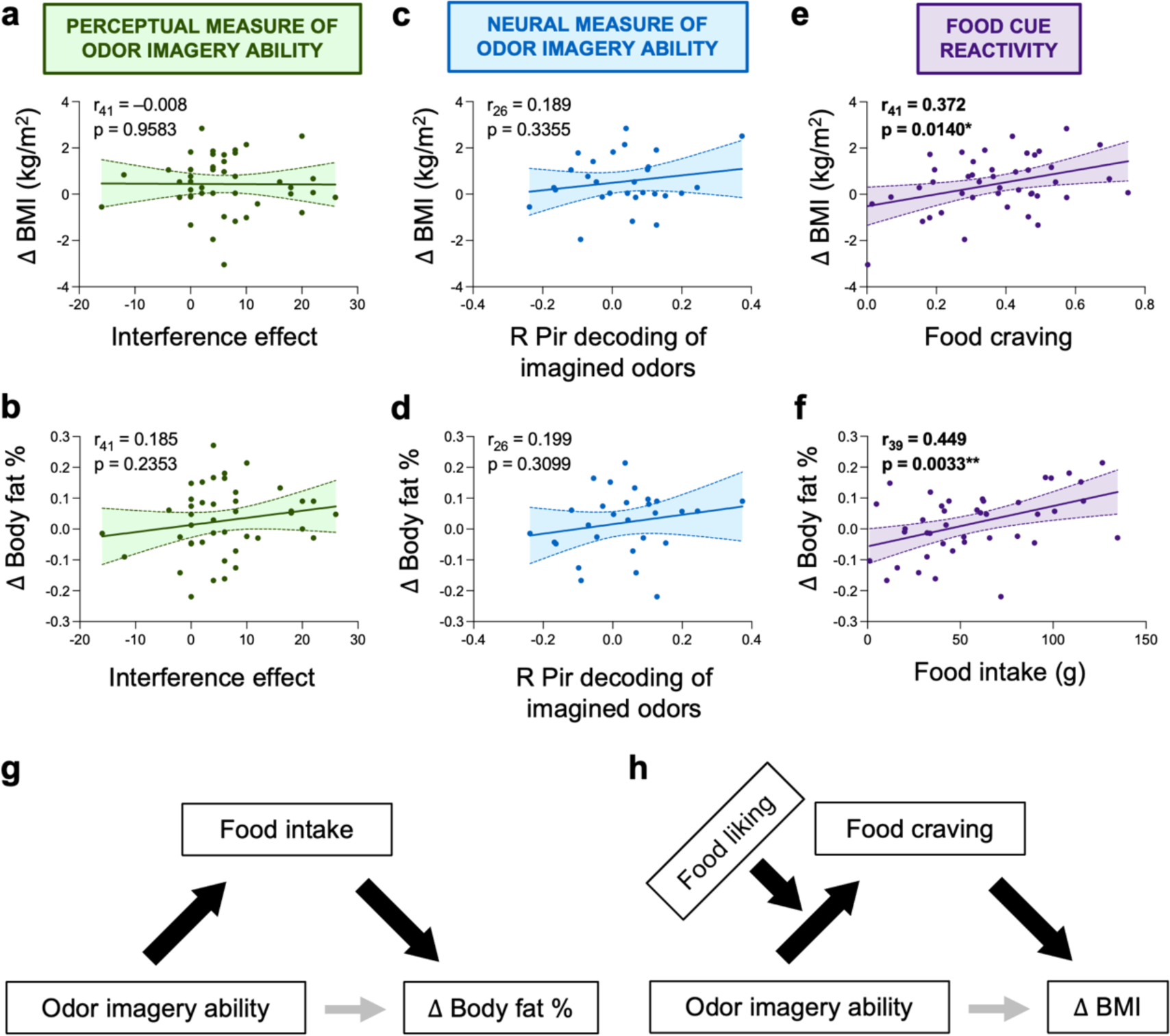
Food Cue Reactivity Mediates the Relationships between Odor Imagery Ability and Changes in BMI and Body Fat Percentage. (a–b) The perceptual measure of odor imagery ability did not correlate with change in BMI (a) or body fat percentage (b). (c–d) The neural measure of odor imagery ability did not correlate with change in BMI (c) or body fat percentage (d). (e–f) Food craving positively correlated with change in BMI (e), whereas food intake positively correlated with change in body fat percentage (f). (g–h) Visualizations of the mediation (g) and moderated mediation (h) models. In both, there was no direct effect between odor imagery ability and adiposity change (thin gray arrows), but the indirect effects via food cue reactivity (thick black arrows) were significant, conditional by food liking in g. Panel g corresponds to Models 1 and 2 and panel h corresponds to Model 3 from Table 2. Fitted scatterplots depict single participants and the 95% CI. R, right; Pir, piriform. *p < 0.05; **p < 0.01.

Given the lack of significant direct effects between odor imagery ability and change in adiposity, we tested for indirect effects via food cue reactivity across three models. The results are summarized in Table 2. Consistent with our hypotheses, cue-potentiated food intake mediated the associations between both the perceptual (Model 1) and neural (Model 2) measures of odor imagery ability and change in body fat percentage (Fig. 6g). In Model 3, we tested whether food craving mediated the relationship between the perceptual measure of odor imagery ability and change in BMI, though here we used moderated mediation to account for the effect of liking on the association between odor imagery ability and craving (Fig. 6h).

**Table 2.** Mediation and Moderated Mediation Models.

| # | Model Path/Effect: Predictor → Outcome | $\beta$ | SE | LL | UL |
| --- | --- | --- | --- | --- | --- |
| 1 | **a: Perceptual measure of odor imagery ability → Food intake** | **0.350** | **0.133** | **0.081** | **0.620** |
| 1 | **b: Food intake → $\Delta$ Body fat %** | **0.505** | **0.182** | **0.136** | **0.874** |
| 1 | c' Direct: Perceptual measure of odor imagery ability → $\Delta$ Body fat % | 0.019 | 0.160 | -0.306 | 0.345 |
| 1 | **a × b Indirect: Perceptual measure of odor imagery ability → Food intake → $\Delta$ Body fat %** | **0.177** | **0.091** | **0.031** | **0.382** |
| 2 | **a: Neural measure of odor imagery ability → Food intake** | **0.407** | **0.166** | **0.063** | **0.751** |
| 2 | **b: Food intake → $\Delta$ Body fat %** | **0.722** | **0.221** | **0.264** | **1.181** |
| 2 | c' Direct: Neural measure of odor imagery ability → $\Delta$ Body fat % | -0.120 | 0.198 | -0.530 | 0.290 |
| 2 | **a × b Indirect: Neural measure of odor imagery ability → Food intake → $\Delta$ Body fat %** | **0.294** | **0.160** | **0.022** | **0.647** |
| 3 | a: Perceptual measure of odor imagery ability → Food craving | -0.180 | 0.133 | -0.449 | 0.090 |
| 3 | **Moderation: Perceptual measure of odor imagery ability × Food liking → Food craving** | **0.368** | **0.112** | **0.141** | **0.595** |
| 3 | **b: Food craving → $\Delta$ BMI** | **0.439** | **0.174** | **0.087** | **0.790** |
| 3 | c' Direct: Perceptual measure of odor imagery ability → $\Delta$ BMI | -0.082 | 0.154 | -0.393 | 0.230 |
| 3 | Conditional a × b indirect (Low food liking): Perceptual measure of odor imagery ability → Food craving → $\Delta$ BMI | -0.079 | 0.103 | -0.347 | 0.055 |
| 3 | Conditional a × b indirect (Moderate food liking): Perceptual measure of odor imagery ability → Food craving → $\Delta$ BMI | 0.033 | 0.103 | -0.347 | 0.158 |
| 3 | **Conditional a × b indirect (High food liking): Perceptual measure of odor imagery ability → Food craving → $\Delta$ BMI** | **0.184** | **0.109** | **0.011** | **0.434** |
| 3 | **Index of moderated mediation: Perceptual measure of odor imagery ability × Food liking → Food craving → $\Delta$ BMI** | **0.161** | **0.104** | **0.007** | **0.411** |

Specifically, food liking was included as a moderator of the a-path. The index of moderated mediation – indicating whether the strength of the indirect effect between odor imagery ability and change in BMI via food craving depended on the level of food liking – was significant (Table 2). In other words, better odor imagery ability resulted in greater changes in BMI through heightened food craving, but only in individuals who liked such high-fat, high-sugar foods. Taken together, these mediation and moderated mediation models provide evidence that odor imagery ability drives variation in food cue reactivity strength, which in turn influences risk for increased adiposity.

## DISCUSSION

It is well established that food cue reactivity including craving is associated with weight gain susceptibility^5^, but the mechanisms underlying this relationship are poorly understood. The current study was motivated by the proposed role for mental imagery in craving intensity^6^ and the existence of significant variation in odor imagery ability^18^ that is positively associated with BMI^20^. These previous observations led us to hypothesize a role for olfactory imagery in driving food cue reactivity strength to promote weight gain. Our results support this hypothesis by demonstrating an indirect link between odor imagery ability and one-year change in adiposity via food cue reactivity. We also show that this effect is selective to imagined odors, as it does not generalize to perceptually experienced odors or to visual imagery.

Mental imagery involves “top-down” reactivation of sensory circuits^35–42^ and is thought to help optimize adaptive behavior through simulations of future actions based on past experiences^43^. In the context of ingestive behavior, food choice depends upon a complex integration of internal and external signals^44^. Imagining what to eat may contribute to food decisions by enabling simulations of the predicted sensory pleasure and eventual nutritive value of eating a potential energy source relative to the current homeostatic state of the organism (e.g., hungry or sated). Thus, imagery facilitates the weighing of the costs and benefits that determine decisions. Indeed, recent preclinical work demonstrates that food odor exposure stimulates lipid metabolism but only in fasted animals with functioning olfactory memory^45^.

Perhaps olfactory memory – a key component of imagery – has the same effect on preparing the body for anticipated intake in humans (and thereby enhancing motivation for food). Our findings suggest that in an environment laden with food cues, the ability to form vivid mental images of the smell and flavor of foods promotes overeating.

In our study, olfactory imagery ability was assessed by multiple measures that correlated with each other and therefore support construct validity. The association between the perceptual measure (i.e., the interference effect) and the neural measure (i.e., right piriform decoding of imagined odors) was particularly strong (Fig. 4f). This suggests that piriform quality coding is critical for, and contributes to the variability in odor imagery ability that has been reported by a number of independent studies^15–19^. Notably, we restricted our correlation analyses to the right piriform cortex because decoding of real odors in this region was significantly greater than both chance level and decoding in the left piriform cortex. This finding is in accordance with evidence for right hemispheric dominance over olfactory processing in general^46, 47^, odor memory^48–50^, and the decoding of real odor quality^51, 52^.

In contrast to our prediction, the crossmodal decoding was not above chance level, meaning that the patterns generated by imagining the odors could not be decoded using the patterns generated by the actual perception of those same odors. One possible explanation for this finding is that the imagined odors only reactivate odor identity while the real odors reactivate odor identity plus the coding of the physiochemical odorant properties occurring across separate subpopulations of piriform cortex neurons^53^. Similar distinctions are observed between imagined and actual coding in other sensory modalities. For example, the neural substrates for the decoding of place memories are immediately anterior to those for real-time scene perception^54^. Likewise, visual imagery engages only a subset of regions contributing to visual perception^55^.

Decoding studies at higher magnetic field strength with smaller voxel sizes would be helpful in testing this possibility.

Our univariate results also align with prior work in olfactory imagery demonstrating responses in the piriform olfactory cortex while imagining odors versus smelling clean air^22, 56, 57^. It is important to note that in the current study, as well as in prior work, the perceptual measure of odor imagery ability (i.e., the interference effect) did not correspond to the magnitude of piriform responses to imagined odors. By contrast, we observed a strong association between the interference effect and imagined odor quality decoding in the right piriform cortex. This suggests that it is these quality codes that underlie imagery, with univariate responses likely including other factors such as sniffing, attention, and associative learning^58^. Accordingly, we ensured that sniffing did not impact our perceptual or neural measures of odor imagery ability. This is critical because sniffing induces piriform activity^59^ and is necessary for the generation of vivid odor imagery^60, 61^. Individuals with better odor imagery ability take larger sniffs while imagining pleasant versus unpleasant smells, a modulatory pattern that is not seen in poor odor imagers^62^. Here we specifically selected pleasant odors to minimize potential sniffing differences.

With respect to attention, the frontal piriform cortex and olfactory tubercle respond preferentially to attended compared to unattended sniffs^63, 64^. In the current study, we instructed participants to sniff in each trial – irrespective of smelling or imagining – prompted by an auditory cue to equate attentional demands. Finally, although visual cues that have been associated with specific odors are capable of evoking piriform^65^ and olfactory bulb^66^ responses, we found that the interference effect correlated with self-reported odor and flavor, but not visual, imagery and that visual imagery did not interfere with detecting an incongruent odor.

Collectively, these data support the conclusion that imagined odor-evoked quality codes in piriform cortex underlie variation in imagery ability rather than non-specific effects such as sniffing, attention, or sensory associative learning.

The principal finding in the current study is that the generation of distinguishable imagined odor quality codes in the piriform cortex correlates not only with imagery ability, but also with measures of food cue reactivity that in turn predict change in adiposity. We therefore propose that better odor imagery leads to stronger food craving and greater intake that promotes obesity risk (Fig. 1c). Accordingly, our mediation analyses revealed that better odor imagery ability does not directly lead to larger adiposity change, but rather that it exerts an influence via a food cue reactivity-dependent mechanism. These results back the Elaborated Intrusion Theory of Desire, which posits that effortful cognitive elaboration of food properties through imagery intensifies cravings^6^. They also corroborate studies showing that vivid sensory imagery is linked to a strong desire for foods, drugs, and alcohol^7–13, 67^. However, they are the first to isolate odor-specific imagery as the critical contributor to food cue reactivity.

The current findings are also relevant to the ongoing work linking olfactory function with risk for weight gain. Although many associations have been reported, the direction is not consistent. Reports for positive^68–73^, negative^74–81^, or no^82, 83^ relationship between olfaction and food intake, current BMI, or weight change have been made. Here we found that olfactory function – defined either as detection thresholds or as piriform decoding of actual odor quality – was unrelated to any measure of odor imagery ability, food cue reactivity, or adiposity change. The same was true for suprathreshold perceptual ratings of odor intensity, familiarity, liking, and edibility. Thus, our results suggest that olfactory simulations or imagery may drive the relationship rather than olfactory coding or perception *per se*, which could account for the inconsistencies that have been noted previously.

For example, imagining what to eat may allow an organism to test the impact of available energy sources on their perceived hunger, pleasure, or mood before selecting which item to consume. It is well known that olfaction is tightly coupled to emotional valence^84^, and odor imagery ability positively correlates with the experience and processing of emotion^16, 85^. The presence of reward-related cues during motor imagery enhances neural activity in and functional connectivity between the motor cortex and ventral striatum, giving rise to a mechanism by which the imagery may become motivationally salient enough to yield action^86^.

While we did not measure emotional reactivity or salience in the current study, we did quantify food liking and observed a moderating effect of this variable. Specifically, we found that better odor imagery ability corresponds to more intense food cravings and larger weight gain in individuals who exhibit strong liking of the high-fat/high-sugar foods that we tested. It is therefore likely that odor imagery interacts with pleasure to invigorate food cravings.

Another possibility is that odor imagery bridges current states with future simulations in guiding food choice through the involvement of the SMA and insula. The SMA is linked to motor planning^87, 88^, while the insula plays a role in interoception and the prediction of bodily states^89–91^. For instance, food cues elicit transient activity across populations of insular neurons that mimics future metabolic conditions^92^ and is necessary for driving food seeking behaviors^93^ in mice. In humans, the SMA and insula are not only consistently activated during mental imagery^22, 94–103^, but also during food and drug craving^14, 104–109^.

Here we identified extensive clusters of activity in the insula preferentially responding to imagining odors versus clean air. Moreover, the left SMA was the only whole-brain-corrected region exhibiting stronger activity while imagining versus smelling odors. These results are in line with the responsivity of the insula and SMA to imagined versus perceived odors in prior work^22, 110^. We also found that the pattern responses predicted by the recently developed Neurobiological Craving Signature^29^, which has positively weighted voxels in both the insula and SMA (among other regions), were greater while imagining a food versus a nonfood odor. These data lead to the hypothesis that information from the coding of imagined odors in the piriform cortex is relayed to the insula and SMA to simulate future interoceptive states and food decisions, promoting the cravings and physiological changes (e.g., ghrelin release) that trigger subsequent consumption. The direct links between the piriform cortex, insula, and SMA and the general validity of this model warrant future testing.

Imagery requires memory systems to pull from past experiences in simulating the future. Though we screened participants for self-reported cognitive deficits or memory loss that could impact mental imagery, we did not explicitly measure memory capacity in the current study.

Impaired memory and hippocampal function are hallmark characteristics in the development of obesity^111^. A recent study reported that despite showing disrupted memory for non-food items, individuals with obesity outperform their lean counterparts in memory for food items^112^.

Importantly, memory was not associated with the perceived vividness of imagining scenes corresponding to the food or non-food cues in that study^112^. The indirect effect of odor imagery ability on change in BMI that we observed here is therefore unlikely linked to memory function, though this should be tested in the context of olfactory imagery in future work. As obesity-related memory alterations may particularly stem from poor diet^113, 114^, the subsequent question emerges of whether frequent consumption of high-fat/high-sugar foods impacts odor imagery ability. Studies do suggest that experience can improve imagery ability, with expert chefs^115^, perfumers^116^, and sommeliers^117^ exhibiting more vivid odor imagery or functional reorganization in its neural correlates. Here we found no relationship between odor imagery ability and typical consumption of energy-dense foods measured with the Dietary Fat and Free Sugar Short Questionnaire^118^ or following the craving or bogus taste test measures. However, future studies should examine the prospective effects of dietary manipulations on odor and flavor imagery ability and, in turn, on food cue reactivity and obesity risk. Food cue reactivity across a wider range of items including nutritional foods should also be considered since there is evidence for increased attention and memory^119^ or selection and intake^31^ of healthy foods following manipulations to sensory appeal or training in cognitive regulation strategies, respectively.

Finally, we note that the current findings have relevance for obesity treatment. One of the leading behavioral strategies for decreasing food cue reactivity is cue exposure therapy (CET)^2^. In CET, patients are trained to refrain from eating their most desired foods during exposure in a controlled setting, thereby extinguishing the learned associations between food cues and consumption. While CET is effective for exposed cues, reductions in food cue reactivity including intake do not generalize to unexposed foods^120, 121^, limiting the potential for weight loss. Our study demonstrates that odor imagery could serve as a novel behavioral therapy target with the VOIQ as a simple tool for screening susceptible individuals. Cognitive tasks that compete with odor imagery may be particularly fruitful in disrupting food cue reactivity^122^. For instance, prior research has shown that imagining the memory of a time that a snack was avoided or thinking about the future consequences of consumption can help to reduce intake in the moment^123, 124^. We propose that imagery in the same sensory modality, such as imagining a nonfood odor or one that is disliked, may prove especially successful in limiting the capacity for flavor imagery to strengthen food cravings.

### Concluding Remarks

In conclusion, the results of our study highlight a role for odor imagery ability in obesity risk via food cue reactivity and point to coding in the piriform primary olfactory cortex as the neural substrate. Future work should explore the extent to which odor imagery helps to integrate internal and external metabolic signals and investigate the efficacy of odor imagery being an additional behavioral target for weight loss therapy.

## MATERIALS AND METHODS

### Participants

A flow diagram depicting the number of individuals at each stage of the study (e.g., eligibility, recruitment, completion, analysis) is provided in Extended Data Fig. 3. Participants were recruited from the local New Haven, CT, USA community and university population via flyer and social media advertisements. Individuals interested in this study or other previous studies in our lab filled out an online form using Qualtrics software (Qualtrics, UT, USA) to indicate initial information such as their sex assigned at birth, age, estimated BMI, drug use, etc. We pre-screened subjects in this database to identify individuals free from known taste or smell dysfunction, dieting behaviors, food restrictions, nicotine or drug use, serious medical conditions including metabolic, neurologic, and psychiatric disorders or medications used to treat these, cognitive deficits or memory loss that could impact mental imagery, and any MRI-contraindications (e.g., being left-handed, pregnant, or having metal in the body). We then assessed for further eligibility with follow-up email questions (e.g., to ensure that these people did not note any new disorders or drug use, recent smell loss due to COVID-19, or intent to leave the greater New Haven, CT area). To capture similar individuals across a range of BMIs, we used stratification to minimize differences in sex, race, ethnicity, age, and household income among individuals recruited into 2 BMI groups (low BMI < 25 and high BMI ≥ 25 kg/m^2^).

For the perceptual measure of odor imagery ability, 36 participants completed all imagery conditions based on a power analysis performed in G*Power version 3.1.9.6^125, 126^ to replicate the interference effect (d = 0.722) from the prior task validation^21^ in the low and high BMI groups (n = 18 each) at 0.80 power (alpha = 0.05, two-tailed test, two dependent means). Twelve additional participants were then recruited to complete only the odor imagery condition and all other study measures (with one excluded from scanning due to extreme claustrophobia). This was sufficient to achieve 0.80 power (n = 42, alpha = 0.05, two-tailed test, bivariate normal model) for the effect observed between self-reported odor imagery ability and obesity risk (r = 0.42) in previous work^20^. Data from three participants were removed due to an inability to obtain proper odor thresholds such that their detection accuracies fell below chance level (less than 50% correct responses). Participant characteristics of the final sample (N = 45) by BMI group are provided in Supplementary Table 8. All individuals provided written informed consent, and the study procedures were approved by the Yale Human Investigations Committee (Institutional Review Board Protocol # 0405026766). The study was also preregistered (AsPredicted.org #56278).

### Stimuli

Odors included “phenylethyl alcohol white extra” (rose, #001059147) and “cookie dough” (cookie, #10610208) from International Flavors and Fragrances (New York, NY, USA) diluted in food-grade propylene glycol. The bogus taste test consisted of eight “Grandma’s Homestyle Chocolate Chip Cookies” broken into bite-sized pieces across two plates (for a total of ∼280g or ∼1360 kcal) presented alongside a 16 fl oz water bottle.

### Experimental Procedures

The study consisted of three behavioral sessions and one fMRI scan at baseline, along with a follow-up session one year later. Full data collection from the first (baseline) to last (follow-up) sessions spanned 10/6/2020–6/3/2022. The fMRI scan was scheduled between 8:00am-1:00pm, and all other sessions took place between 8:00am-8:00pm. We ensured that food craving and intake were assessed between the hours of 11:30am-7:00pm. Individuals were instructed to arrive to all sessions neither hungry nor full, but at least one-hour fasted.

#### Behavioral Sessions

##### Training and Scales

Participants were first trained to make computerized ratings in PsychoPy version 3.0^127^ by practicing with imagined sensations (e.g., the taste of your favorite chocolate) and real stimuli (e.g., the brightness of the ceiling light or the pressure of a weight). Intensity and liking were rated with the vertical category-ratio general Labeled Magnitude Scale (gLMS)^128–130^ and Labeled Hedonic Scale (LHS)^33^, respectively. The gLMS ratings were log base 10 transformed prior to any analyses. All other ratings were made on horizontal visual analog scales (VAS). Familiarity and edibility were rated from “not at all familiar” to “more familiar than anything” and from “not at all” to “more than anything” in response to “how much do you want to eat this?”, respectively. Internal state ratings for hunger, fullness, thirst, anxiety, and need to urinate were made from “not at all [hungry/full/etc.]” to “more [hungry/full/etc.] than anything.” Subjective hunger was calculated as the difference of VAS ratings for hunger – fullness. Participants also practiced one odor run in a mock MRI simulator in the lab.

##### Adiposity

Body weight was measured with an electronic scale and height with a digital stadiometer to calculate BMI. Bioelectric impedance analysis (Seca Medical Body Composition Analyzer mBCA 525, Hamburg, Germany) was used to obtain body fat percentage; values were divided by 21 for females and by 31 for males to adjust for sex.

##### Questionnaires

Participants completed the Vividness of Olfactory Imagery^30^ and Vividness of Visual Imagery^17^ Questionnaires (VOIQ/VVIQ) in which they imagined odors/visual objects across 16 scenarios and rated the vividness of their mental imagery from one “perfectly clear and as vivid as normal smell/vision” to five “no image at all – you only know you are thinking of an odor/object.” Both inventories were reverse scored such that higher sums reflected larger self-reported imagery ability. Participants also did a modified Vividness of Food Imagery Questionnaire (VFIQ)^20^ that was similar to the VOIQ but focused on the ability to imagine external food odors (e.g., of cookies in the oven) and flavors in the mouth (e.g., of eating cookies, which also rely on olfaction). Total weekly metabolic equivalent task-minutes (MET-minutes) from the International Physical Activity Questionnaire (IPAQ)^131^ and total score from an American version of the Dietary Fat and Free Sugar Short Questionnaire (DFS)^118^ were also used to assess habitual exercise and high-fat/high-carbohydrate intake, respectively. MET-minutes for each type of physical activity represent the total minutes dedicated to the activity times the estimated energy expenditure during the activity as a multiple of resting energy expenditure (e.g., vigorous activities count toward a higher MET score than moderate activities).

##### Perceptual Task of Odor Imagery Ability

Detection thresholds for the rose and cookie odors were first determined using a 16-step dilution series (4% odor by volume to 1.22ppm) in a 2-alternative forced-choice staircase procedure^132^. In a within-subjects and counterbalanced design, participants then completed three imagery conditions (odor, visual, and none) of a validated perceptual task^21^. During odor and visual imagery, they were instructed to imagine the smell or sight of one odor type (e.g., rose) while trying to determine which of two samples “smelled stronger.” In matched trials, the two samples contained: (1) the same odor as the imagined type – e.g., rose – at their detection threshold level, and (2) the odorless propylene glycol diluent. In mismatched trials, the two samples were: (1) the incongruent odor – e.g., cookie, and (2) the odorless diluent. In the no imagery condition, odor detection trials were performed in the absence of imagery. The odor and visual imagery conditions contained 25 matched and 25 mismatched trials per odor (100 total), and the no imagery condition consisted of 25 trials per odor (50 total), all counterbalanced for presentation order (i.e., sample one contained the odor in 50% of trials). The interference effect (perceptual measure of odor imagery ability) was calculated by subtracting detection accuracy (% trials correct) in mismatched trials from that in matched trials of the odor imagery condition.

*Food Cue Reactivity.* Cue-induced craving strength was rated in response to 90 palatable food pictures^31^ on a horizontal VAS from “I do not want it at all” to “I crave it more than anything,” and the average was calculated. Items included familiar American snacks and meals, such as pizza and doughnuts. For cue-potentiated intake, participants completed a bogus taste test^32^ in which they were instructed to eat as much as they liked while comparing the sensory properties of two plates of cookies (e.g., which tastes sweeter/saltier, is fresher, or has better quality chocolate). They were not explicitly told that the cookies were identical and that the primary aim was to quantify the grams consumed. Data from two participants were excluded from this measure after eating more than 3 SD above the group mean. Following the food craving and intake paradigms, participants also rated their liking on the LHS^33^ and frequency of consumption in a typical month on a VAS (labels: 1 or less/month, 2/month, 3/month, 1/week, 2/week, 3–4/week, 5–6/week, 1/day, 2 or more/day) for each stimulus.

#### fMRI Session

Participants underwent fMRI scanning while performing a task in an event-related design with six trial types: smell rose, cookie, or clean air; and imagine rose, cookie, or clean air. Each trial began with a 5s auditory cue of “smell” or “imagine” followed by the name of the odor (e.g., “rose”) and the countdown “three, two, one, sniff.” Odor/clean air delivery (3s) was time-locked to sniff onset. Trials were separated by intertrial intervals of 7–17s (mean = 10s). Participants completed 30 trials per run (five of each type) and five runs per scan. Runs were ∼9min long and separated by ∼2min breaks to minimize olfactory habituation.

Stimuli were delivered at concentrations matching individual ratings of moderate intensity on the gLMS with a custom MRI-compatible olfactometer that has been described in detail previously^133^. In brief, the odors and clean air were presented via tubing channels and removed by a vacuum line connected to a NuancePro Gel Nasal Pillow Fit-Pack Model #1105167 nasal mask (Philips Respironics, Murrysville, PA, USA) worn by the subject. This mask was coupled to an anti-viral filter (item #28350, Vitalograph, Lenexa, KS, USA) followed by a pneumotachograph to measure airflow in the nose, which was then attached to a spirometer and amplified with PowerLab 4SP for digital recording at 100 Hz in LabChart version 7 (ADInstruments, Sydney, Australia). Participants completed pre- and post-scan odor and internal state ratings in the MRI bore before and after scanning. These ratings were averaged and the differences of cookie versus rose intensity, familiarity, liking, and edibility were quantified (Extended Data Fig. 4).

fMRI data were acquired with a Siemens 3 Tesla Magnetom Prisma scanner using a 32- channel head coil. Images were collected at an angle of 30° off AC-PC to reduce susceptibility artifacts in olfactory regions. Sagittal T1 anatomical images (repetition time TR = 1900ms, echo time TE = 2.52ms, 176 slices, field of view FOV = 250mm, voxel size = 1×1×1mm) and functional echo-planar images (EPIs) with a multiband BOLD sequence (TR = 2100ms, TE = 40ms, 72 slices, flip angle = 85°, FOV = 192mm, voxel size = 1.5×1.5×1.5mm) were obtained.

#### Follow-Up Session

All but one participant returned to the lab approximately one year later (days elapsed from first to last session: M = 363.17, SD = 7.33, range = 340 – 378) to repeat the adiposity, questionnaire, and food cue reactivity measures. Follow-up data from one participant was excluded after they began a strict diet and lost more than 3 SD above the group mean in weight change from the baseline to follow-up sessions.

### Data Analyses

#### Behavioral Analyses

Pearson correlations, linear regressions, linear mixed effects models, ANOVAs, and Student’s t-tests were performed in MATLAB 2020a (Mathworks, Natick, Massachusetts, USA). Data were plotted in Prism version 9.4.1 (GraphPad Software, San Diego, CA, USA). Mediation and moderated mediation models were tested with bootstrapping (10000 samples, 95% CIs) using the “PROCESS” macro version 4.1^134^ models 4 and 7 implemented in SPSS Statistics version 28 (IBM, Chicago, IL, USA). Significant effects were supported by confidence intervals (CIs) excluding zero within the lower and upper bounds. For test-retest reliability, intraclass correlation coefficient estimates and 95% CIs were calculated in SPSS based on single measure, absolute agreement, 2-way mixed models. All measures showed moderate to good reliability (Supplementary Table 9).

##### Sniff Analyses

The spirometer data were preprocessed and analyzed in MATLAB R2020a. The raw airflow traces were separated by scan run and preprocessed (temporally smoothed with a 500ms moving window, high-pass filtered at a cutoff of 0.02 Hz, and normalized by subtracting the mean and dividing by the SD). Sniff onset for each trial was determined by finding the time of the minimum signal value within a window of ± 0.75s from the auditory cue end. The time (latency) and value (amplitude) of the proximal maximum signal were identified. Sniff offset was defined as the time (duration) at which the signal returned to its original minimum, which was used in quantifying the area under the curve with the trapezoidal method (volume). Finally, peak and mean airflow rates were assessed using derivatives at each signal point indicating the instantaneous rates of change. These parameters were averaged by trial type (e.g., smell rose) for each participant prior to comparison in ANOVAs (Extended Data Fig. 5 and Supplementary Table 2).

#### fMRI Analyses

##### Preprocessing

The fMRI data were preprocessed and analyzed using FSL version 5.0.10 (FMRIB Software Library, Oxford, UK; Jenkinson et al., 2012) and SPM12 (Statistical Parametric Mapping, Wellcome Centre for Human Neuroimaging, London, UK) implemented in MATLAB R2019b. Functional EPIs were realigned to the mean and unwarped using fieldmaps, slice-time corrected, and motion-corrected with the FSL tool MCFLIRT^136^. The anatomical T1 image was coregistered to the mean EPI and spatially normalized to the standard MNI reference with unified segmentation in SPM12. Prior to the univariate analyses, the resulting deformation fields were applied to the EPI images, which were then smoothed with a 3mm full-width-half-maximum Gaussian kernel.

##### First Level Models

General linear models (GLMs) were estimated for each participant and run, separately for the normalized and smoothed EPI data (for univariate analyses) and the non-normalized and non-smoothed EPI data (for decoding analyses). In each, the 6 trial types (smell rose/cookie/clean air and imagine rose/cookie/clean air) were modeled with a canonical hemodynamic response function as events of interest with onsets time-locked to the start of odor/clean air delivery and durations of 3s. The following nuisance regressors were also included: 24 motion parameters (the six SPM realignment parameters for the current volume, six for the preceding volume, plus each of these values squared^137^, the mean signal extracted from the ventricular cerebrospinal fluid computed with *fslmeants*, a matrix of motion-outlier volumes identified using *fsl_motion_outliers* (threshold = 75^th^ percentile plus 2.5 times the interquartile range and/or greater than 1mm of framewise displacement^138^), and the preprocessed sniff trace down-sampled to the scanner temporal resolution with decimation. A 128s high-pass filter was applied to remove low-frequency noise and slow signal drifts.

##### Univariate Analyses

The following contrast images were created at the single-subject level and averaged across the five runs: smell odor (rose + cookie) > smell clean air, imagine odor > imagine clean air, imagine odor > smell clean air, smell odor > imagine odor, and imagine odor > smell odor. Group-level random effects analyses were conducted with one-sample t-tests thresholded at p_uncorrected_ < 0.001 and a cluster size of at least five contiguous voxels. Effects were considered significant at p < 0.05, cluster-level family-wise error corrected across the whole brain. We also regressed the perceptual measure of odor imagery ability (i.e., the interference effect) against whole-brain BOLD responses to imagining odors > imagining clean air and imagining odors > smelling clean air. Here we considered whole-brain effects and those significant in the piriform cortex at a peak-level of p < 0.025, family-wise error small-volume corrected for multiple comparisons in our two regions of interest (see below) and subsequently Bonferroni corrected for the two SVC searches. The anatomical labels were determined jointly from the “Atlas of the Human Brain”^139^, an adult maximum probability atlas prepared with SPM12 (www.brain-development.org)^140-142^, and the Automated Anatomical Labeling Atlas 3^143^.

*Decoding Analyses.* The ROIs for the decoding analyses included the left and right piriform cortices independently created from the Neurosynth^144^ meta-analytic functional map for the term “olfactory” (74 studies with 2021 activations, downloaded 9/15/2021). Activations from this map were restricted to a threshold of z = 6 to ensure separability of the piriform clusters from other nearby regions (e.g., the insula). The ROIs were converted from MNI space to each subject’s native EPI space (voxel size = 1.5×1.5×1.5mm), resulting in clusters of 190 and 111 voxels for the left and right piriform, respectively.

MVPA was performed using The Decoding Toolbox^145^ implemented in SPM12. For the first decoding method (SVM classification), separate voxel-wise patterns were created for smelling and imagining the rose and cookie odors by extracting the parameter estimates from the first level GLMs and subtracting the mean activity across the conditions in each run. Feature selection was used to identify the top class-discriminative voxels in each ROI with an ANOVA, restricted to the number of voxels in each ROI maximally available for all subjects. An SVM from the Library for Support Vector Machines (LIBSVM) package^146^ was trained to decode rose versus cookie using patterns of BOLD activation for smelling the odors in four of five scan runs. The SVM was then tested for its accuracy to predict these odor categories from the patterns in the left-out run. These steps were repeated for training and tested on the imagined odor patterns, and for training on smelled and testing on imagined (and vice versa, averaged for the crossmodal condition). SVM accuracies were compared to chance (50%) in one-sample t-tests to assess group-level significance. SVM accuracies for the decoding of real odors in the left versus right piriform cortex were also directly compared with a paired-samples t-test to assess the laterality of the effect.

For the second decoding method (split-half voxel correlations), the first BOLD run was treated as an odor localizer, which resulted in an equivalent number of even and odd runs remaining for decoding (2 each). The voxels for each subject and ROI were functionally ranked according to their t values in the contrast of smelling odor > smelling clean air from the localizer. Again, the N-most odor-active voxels maximally available for all subjects were selected. The split-half voxel correlations were then analyzed for the within-odor (e.g., smelling rose in even runs versus smelling rose in odd runs) minus the between-odor (e.g., smelling rose in even runs versus smelling cookie in odd runs) fMRI patterns in each ROI. In line with our SVM analyses, we performed separate tests for real, imagined, and crossmodal odors. The resulting correlation values were Fisher’s Z transformed and compared to zero in one-sample t-tests to assess group-level significance. They were also tested in correlations against the perceptual and self-report measures of odor imagery ability. The latter analyses were performed in all individuals (n = 44) and separately restricted to those with discriminable neural patterns for actual odors, defined as within-odor minus between-odor voxel correlation Z-values > 0 (n = 30).

##### Testing the Neurobiological Craving Signature

The NCS is a recently developed neuromarker or brain signature^34^ of craving^29^ that predicts the intensity of drug and food craving with good accuracy. To assess the responses of the NCS-food pattern (a pattern that was trained on visual food cues only), we computed the matrix dot product between this NCS-food weight map and each participant’s L2-normed contrast images for the six conditions: smell cookie/rose/clean air and imagine cookie/rose/clean air. The matrix dot product provides one scalar value (a ‘pattern response’ value) per participant and contrast image that describes the similarity of the image to the weight map and the predicted level of food craving. Greater responses of the NCS-food weight map indicate greater similarity to the craving map and higher predicted levels of food craving. Pattern response values were statistically compared with an ANOVA and paired t-tests for planned comparisons. All weight maps and code to apply the NCS are publicly available at: https://github.com/canlab/Neuroimaging_Pattern_Masks/tree/master/Multivariate_signature_patterns/2022_Koban_NCS_Craving.

## DATA AVAILABILITY

The raw MRI data and sniff airflow traces can be downloaded from the OpenNEURO repository under accession number ds004327: https://openneuro.org/datasets/ds004327. Statistical maps of the human brain will be made available on the NeuroVault repository.

## ACKNOWLEDGEMENTS

This work was supported by the National Science Foundation Graduate Research Fellowship under Grant No. 2139841 (E.E.P.), the National Institute of Diabetes and Digestive and Kidney Diseases of the National Institutes of Health under Award No. F31DK130556 (E.E.P.), and the Modern Diet and Physiology Research Center (D.M.S.). The content is solely the responsibility of the authors and does not necessarily represent the official views of the National Science Foundation or the National Institutes of Health. We would like to thank Jason Avery for advice on the fMRI decoding methods; Bojana Kuzmanovic for guidance on the fMRI preprocessing pipeline; James Howard for example code to perform the sniffing analyses; Thomas Hummel, Johan Lundström, Joel Mainland, and Paul Wise for their thoughts in troubleshooting the odor detection threshold testing; Alain Dagher, Ralph DiLeone, and Barry Green for their helpful suggestions to the project design; and Karen Martin for MR technical assistance.

## AUTHOR CONTRIBUTIONS

Conceptualization, E.E.P. and D.M.S.; Methodology, E.E.P., X.S.D., J.D., M.J-G., J.T., Z.H., M.G.V., H.K., and D.M.S.; Formal Analysis, E.E.P., X.S.D., and L.K.; Investigation, E.E.P. and J.T.; Resources, X.S.D., J.D., M.J-G., Z.H., M.G.V., T.W., H.K., and D.M.S.; Data Curation, E.E.P.; Writing – Original Draft, E.E.P. and D.M.S.; Writing – Review & Editing, X.S.D., J.D., M.J-G., J.T., Z.H., M.G.V., L.K., T.W., and H.K.; Visualization, E.E.P.; Supervision, X.S.D., H.K., and D.M.S.; Funding Acquisition, E.E.P. and D.M.S.

## EXTENDED DATA FIGURES

**Extended Data Fig. 1:**
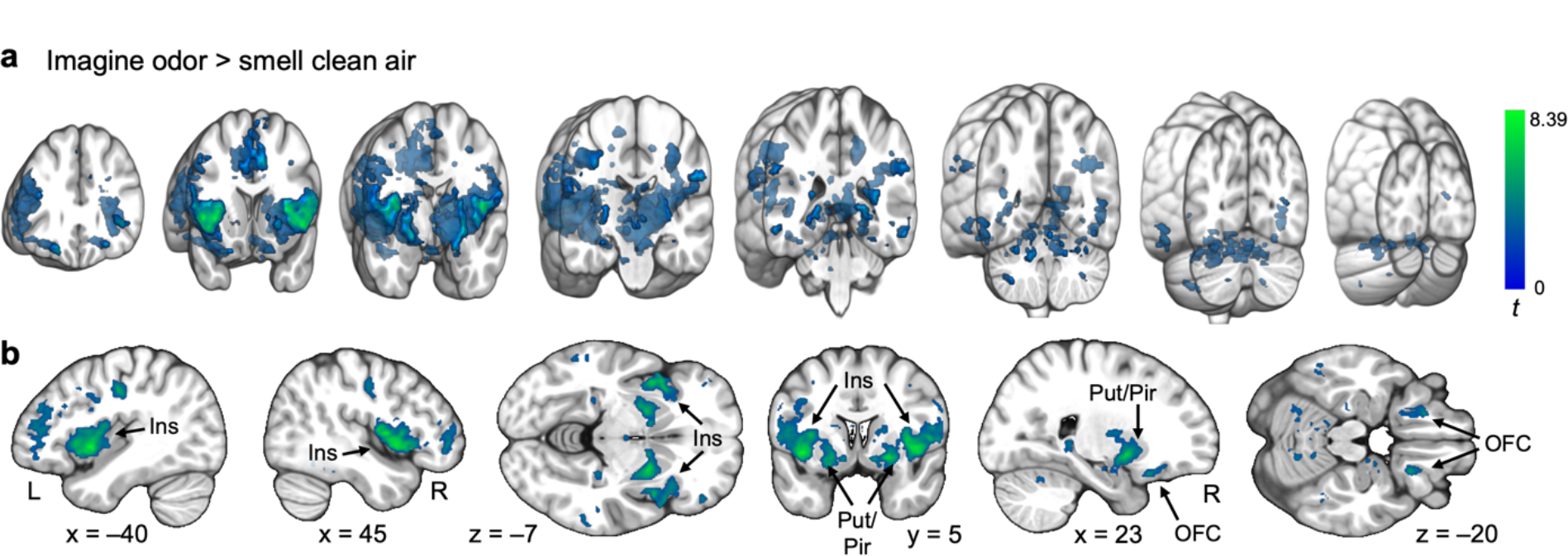
Univariate fMRI Activity to Imagining Odors > Smelling Clean Air. (a) 3D coronal sections (18mm thick) evenly spanning y = 56 to –88mm depict significant BOLD responses to imagining odors > smelling clean air in the bilateral insula, putamen extending into the piriform cortices, pallidum, and orbitofrontal, middle frontal, and precentral gyri, among other regions. (b) Important areas of activation for imagining odors > smelling clean air are highlighted with custom coordinates (see Supplementary Table 4). Brain sections show the SPM *t*-map (*p*uncorrected < 0.005, clusters of at least 5 voxels) overlaid onto an anatomical template in MNI coordinates for illustrative purposes. Color bars depict *t* values. L, left; R, right; Ins, insula; OFC, orbitofrontal cortex; Pir, piriform cortex; Put, putamen.

**Extended Data Fig. 2:**
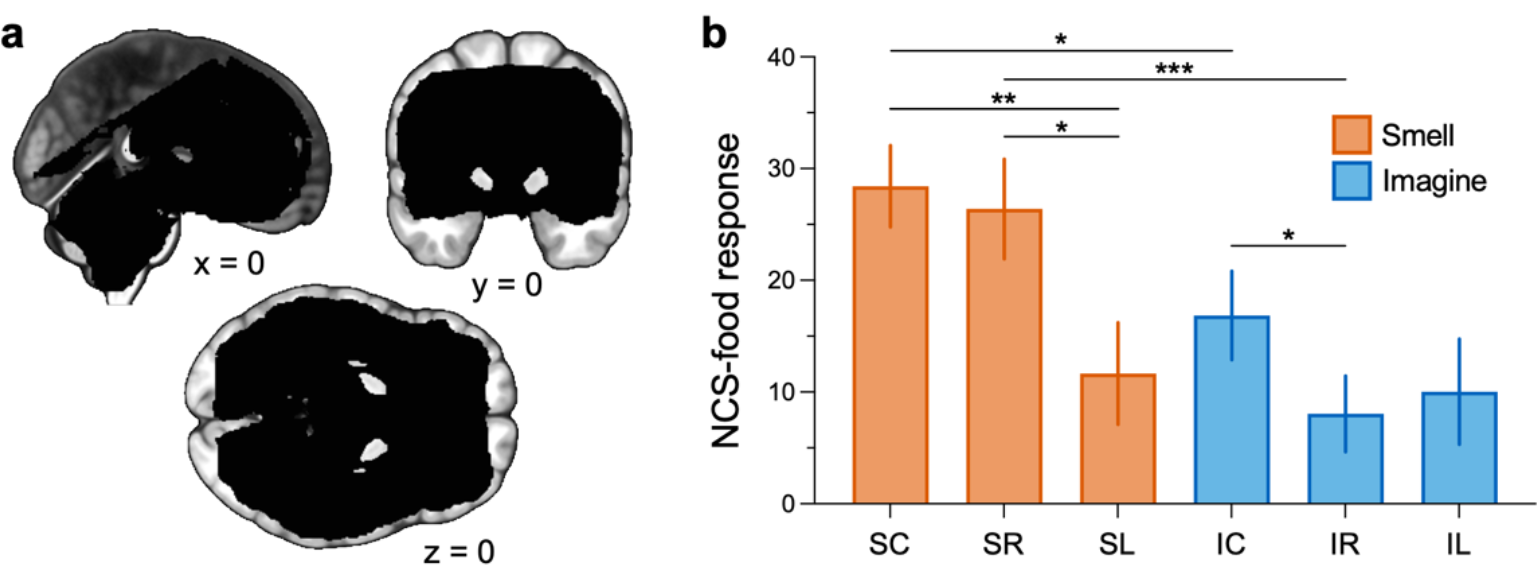
Imagining a Food Odor Elicits Greater Neurobiological Craving Signature Activation than Imagining a Nonfood Odor. (a) Mask constructed from the intersection of EPI scan windows for all participants (black) overlaid onto an anatomical template in MNI coordinates to depict the fMRI signal coverage. (b) The effects of smelling and imagining cookie and rose odors and clean air on pattern responses of the recently developed Neurobiological Craving Signature (NCS) from independent work^29^. S, smell; I, imagine; C, cookie; R, rose; L, clean air. Post-hoc comparisons: *p < 0.01, **p < 0.001, ***p < 0.0001.

**Extended Data Fig. 3:**
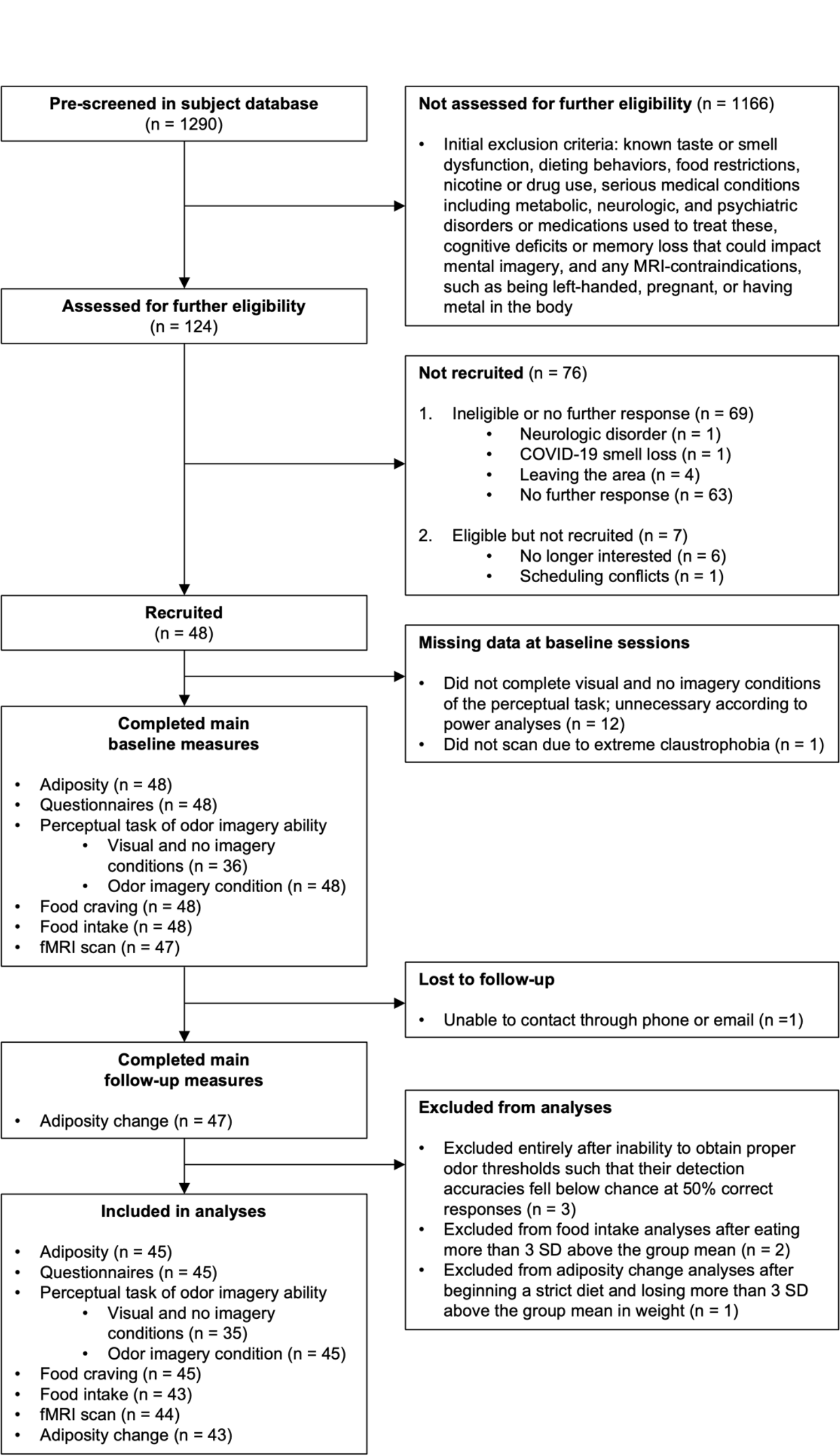
Participant Flow Diagram. Flow diagram depicting the number of individuals at each stage of the study.

**Extended Data Fig. 4:**
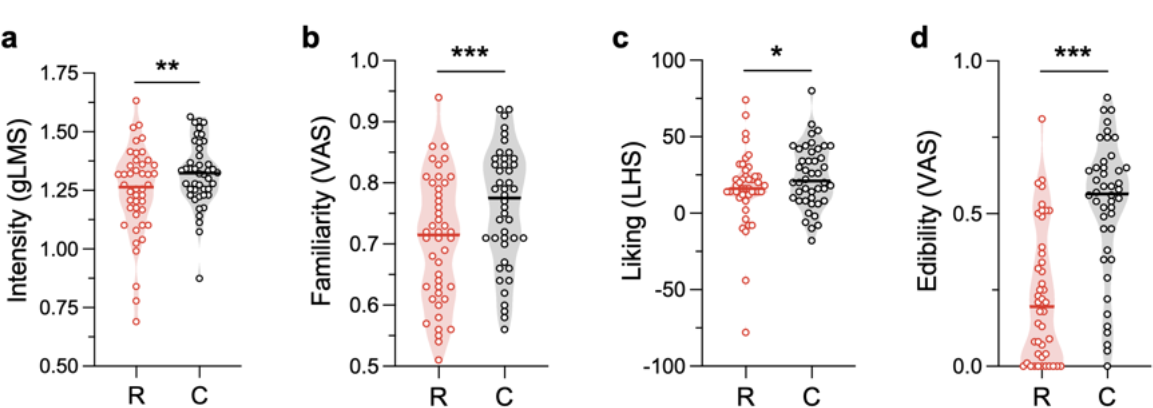
Odor Rating Comparisons for Rose Versus Cookie. (a–d) The cookie odor was rated to be significantly more intense (a), familiar (b), liked (c), and edible (d) than the rose odor. However, the cookie minus rose odor ratings were not correlated with any measure of odor imagery ability (Supplementary Table 1). Truncated violin plots depict single participants with shading to represent the density of the points around the median line. R, rose; C, cookie; gLMS, general Labeled Magnitude Scale^128–130;^ VAS, visual analog scale; LHS, Labeled Hedonic Scale^33^. *p < 0.05; **p < 0.01; ***p < 0.0001.

**Extended Data Fig. 5:**
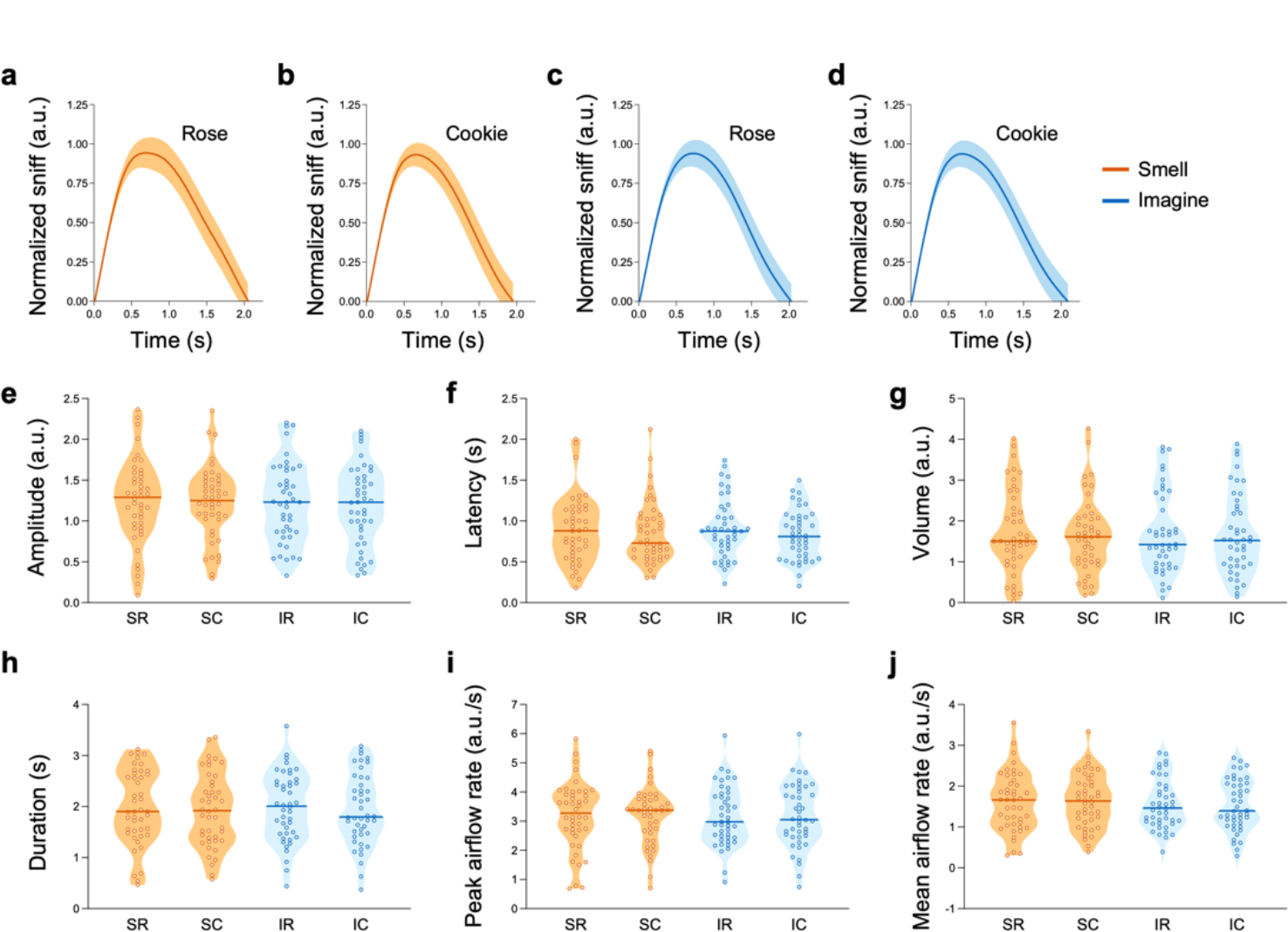
Sniff Parameters for Smelling and Imagining the Rose and Cookie Odors. (a–d) Normalized sniff traces (M ± SEM) for smelling the rose (a) and cookie (b) odors and imagining the rose (c) and cookie (d) odors. (e–j) Sniff amplitude (e), latency (f), volume (g), duration (h), peak airflow rate (i), and mean airflow rate (j) while smelling and imagining the rose and cookie odors. Differences in the sniff parameters for imagining the cookie minus rose odor were not correlated with any measure of odor imagery ability (Supplementary Table 1). ANOVAs also revealed no main effects or interactions of modality (smell/imagine), odor (rose/cookie), or the perceptual measure of odor imagery ability (the interference effect) on any sniff parameter (Supplementary Table 2). Truncated violin plots depict single participants with shading to represent the density of the points around the median line. S, smell; I, imagine; R, rose; C, cookie; a.u., arbitrary units.

