## Supplementary Materials for "Odor imagery but not perception drives risk for food cue reactivity and increased adiposity"

**Supplementary Table 1. Correlations Comparing the Odor Imagery Ability Measures with Demographics, Current Adiposity, Olfactory Function, Odor Ratings, Sniff Parameters, Hunger, and Consumption of Unhealthy Foods**

| Variable (Units) | Self-Reported (VOIQ Score) | | Perceptual (Interference Effect) | | Neural (Right Piriform Decoding of Imagined Odors) | |
|---|---|---|---|---|---|---|
| | r | p | r | p | r | p |
| Sex | <0.001 | 0.9958 | 0.021 | 0.8916 | 0.206 | 0.2753 |
| Age (yr) | 0.176 | 0.2487 | 0.176 | 0.2477 | 0.215 | 0.2538 |
| Household Income | –0.045 | 0.7692 | 0.090 | 0.5588 | 0.181 | 0.3394 |
| BMI (kg/m$^2$) | 0.225 | 0.1381 | 0.011 | 0.9414 | 0.021 | 0.9107 |
| Body Fat % | 0.232 | 0.1257 | 0.115 | 0.4533 | 0.125 | 0.5106 |
| Rose Odor Threshold | –0.171 | 0.2611 | –0.058 | 0.7061 | –0.129 | 0.4986 |
| Cookie Odor Threshold | –0.280 | 0.0630 | 0.199 | 0.1890 | –0.220 | 0.2429 |
| Odor Liking | 0.163 | 0.2893 | 0.179 | 0.2446 | 0.124 | 0.5132 |
| Odor Edibility | 0.047 | 0.7598 | 0.173 | 0.2608 | 0.144 | 0.4485 |
| Odor Intensity | –0.113 | 0.4671 | –0.081 | 0.6028 | –0.223 | 0.2365 |
| Odor Familiarity | –0.166 | 0.2828 | –0.2805 | 0.0651 | –0.162 | 0.3921 |
| Sniff Amplitude | –0.091 | 0.5555 | 0.022 | 0.8857 | –0.074 | 0.6966 |
| Sniff Latency | –0.296 | 0.0508 | –0.092 | 0.5522 | –0.260 | 0.1646 |
| Sniff Volume | –0.227 | 0.1389 | –0.012 | 0.9399 | –0.138 | 0.4667 |
| Sniff Duration | –0.205 | 0.1811 | –0.088 | 0.5685 | –0.183 | 0.3319 |
| Sniff Peak Airflow Rate | 0.052 | 0.7391 | 0.100 | 0.5200 | –0.303 | 0.1040 |
| Sniff Mean Airflow Rate | 0.040 | 0.7954 | 0.104 | 0.5003 | 0.081 | 0.6694 |
| Hunger | –0.099 | 0.5160 | –0.121 | 0.4305 | –0.286 | 0.1251 |
| DFS Score | –0.069 | 0.6511 | –0.033 | 0.8312 | 0.018 | 0.9237 |
| Frequency of Consumption for Craving Stimuli | 0.011 | 0.9515 | –0.013 | 0.9312 | –0.092 | 0.6272 |
| Frequency of Consumption for Intake Stimuli | 0.183 | 0.2286 | 0.198 | 0.1933 | 0.090 | 0.6380 |
| Right Piriform Decoding of Actual Odors | 0.039 | 0.8362 | –0.185 | 0.3290 | 0.038 | 0.8403 |

The odor ratings (liking, edibility, intensity, and familiarity) were quantified as the difference of cookie minus rose. The sniff parameters (amplitude, latency, volume, duration, peak airflow rate, and mean airflow rate) were quantified as the difference in imagine cookie minus imagine rose trials. See also Extended Data Figs. 4 and 5 for further comparisons of the odor ratings and sniff parameters by trial type and Supplementary Table 2 for main effects and interactions of modality (smell/imagine), odor type, and the perceptual measure of odor imagery ability on the sniff parameters. DFS, Dietary Fat and Free Sugar Short Questionnaire[118], American version; VOIQ, Vividness of Olfactory Imagery Questionnaire[30].

**Supplementary Table 2. ANOVAs Comparing the Sniff Parameters by Modality (Smell/Imagine), Odor Type (Rose/Cookie), and the Perceptual Measure of Odor Imagery Ability (i.e., the Interference Effect)**

| ANOVA Main Effects and Interactions | Amplitude | | Latency | | Volume | | Duration | | Peak Airflow Rate | | Mean Airflow Rate | |
|---|---|---|---|---|---|---|---|---|---|---|---|---|
| | $F_{1,168}$ | p | $F_{1,168}$ | p | $F_{1,168}$ | p | $F_{1,168}$ | p | $F_{1,168}$ | p | $F_{1,168}$ | p |
| Modality | 0.092 | 0.7623 | 0.938 | 0.3343 | 0.006 | 0.9401 | 0.373 | 0.5424 | 0.081 | 0.7760 | 0.442 | 0.5071 |
| Odor Type | 0.021 | 0.8838 | 0.681 | 0.4103 | 0.661 | 9.4174 | 0.121 | 0.7284 | 0.012 | 0.9122 | 0.017 | 0.8978 |
| Odor Imagery Ability | 0.191 | 0.6625 | 0.347 | 0.5565 | 0.009 | 0.9233 | 0.244 | 0.6222 | 0.848 | 0.3586 | 0.202 | 0.6534 |
| Modality × Odor Type | 0.332 | 0.5650 | 0.153 | 0.6959 | 0.225 | 0.6355 | 0.0002 | 0.9889 | 0.010 | 0.9198 | 0.148 | 0.7007 |
| Modality × Odor Imagery Ability | 1.882 | 0.1719 | 1.303 | 0.2552 | 1.272 | 0.2609 | 1.162 | 0.2826 | 0.615 | 0.4339 | 0.120 | 0.7291 |
| Odor Type × Odor Imagery Ability | 0.588 | 0.4445 | 0.015 | 0.9035 | 0.594 | 0.4421 | 0.658 | 0.4186 | 0.061 | 0.8048 | 0.384 | 0.5365 |
| Modality × Odor Type × Odor Imagery Ability | 0.413 | 0.5214 | 0.094 | 0.7598 | 0.236 | 0.6276 | 0.025 | 0.8757 | 0.041 | 0.0840 | 0.011 | 0.9175 |

See Extended Data Fig. 5 for comparisons of the sniff parameters by trial type.

**Supplementary Table 3. Brain Regions with Significant Responses to Smelling Odors > Smelling Clean Air**

| Left/Right Label (Brodmann Area) | Size (Voxels) | $p_{FWE}$ | t | MNI | | |
|---|---|---|---|---|---|---|
| | | | | x | y | z |
| **L insula anterior long gyrus** | **3377** | **<0.0001** | **9.700** | **–37.5** | **2** | **–10** |
| L insula posterior short gyrus | | | 9.474 | –36 | –1 | 0.5 |
| L piriform cortex | | | 9.149 | –28.5 | 2 | –17.5 |
| **R amygdala** | **4198** | **<0.0001** | **9.173** | **19.5** | **–2.5** | **–17.5** |
| R insula anterior long gyrus (38) | | | 8.469 | 33 | 6.5 | –17.5 |
| R insula anterior long gyrus | | | 7.622 | 39 | 2 | –7 |
| **R posterior orbitofrontal gyrus (47)** | **301** | **<0.0001** | **6.731** | **21** | **24.5** | **–19** |
| R posterior orbitofrontal gyrus | | | 4.235 | 27 | 30.5 | –14.5 |
| R anterior orbitofrontal gyrus (11) | | | 3.646 | 25.5 | 42.5 | –11.5 |
| **L cerebellum declive** | **291** | **<0.0001** | **6.028** | **–19.5** | **–61** | **–23.5** |
| L cerebellum culmen | | | 4.944 | –30 | –58 | –29.5 |
| L cerebellum declive | | | 3.879 | –25.5 | –68.5 | –23.5 |

| | | | | | | |
|---|---|---|---|---|---|---|
| **L precentral gyrus** | **134** | **0.0001** | **5.797** | **–39** | **–16** | **38** |
| L precentral gyrus | | | 4.116 | –46.5 | –13 | 41 |
| **R middle frontal gyrus** | **486** | **<0.0001** | **5.741** | **42** | **47** | **6.5** |
| R middle frontal gyrus | | | 5.277 | 39 | 41 | 11 |
| R middle frontal gyrus (10) | | | 4.033 | 45 | 47 | 14 |
| **L posterior orbitofrontal gyrus** | **403** | **<0.0001** | **5.705** | **–24** | **30.5** | **–20.5** |
| L posterior orbitofrontal gyrus | | | 4.885 | –31.5 | 33.5 | –11.5 |
| L posterior orbitofrontal gyrus | | | 4.645 | –19.5 | 17 | –22 |
| **L postcentral gyrus** | **151** | **<0.0001** | **5.344** | **–58.5** | **–17.5** | **18.5** |
| **L posterior cingulate gyrus (23)** | **346** | **<0.0001** | **5.339** | **–1.5** | **–25** | **29** |
| R posterior cingulate gyrus | | | 5.308 | 6 | –17.5 | 30.5 |
| Posterior cingulate gyrus | | | 4.718 | 0 | –35.5 | 26 |
| **R thalamus** | **57** | **0.0324** | **5.292** | **4.5** | **–16** | **0.5** |
| R thalamus | | | 3.444 | 12 | –19 | –1 |
| **R postcentral gyrus** | **300** | **<0.0001** | **5.151** | **60** | **–16** | **23** |
| R supramarginal gyrus | | | 4.866 | 60 | –37 | 32 |
| R supramarginal gyrus (40) | | | 4.660 | 63 | –32.5 | 26 |
| **R cerebellum declive** | **130** | **0.0001** | **4.851** | **22.5** | **–61** | **–25** |
| R cerebellum culmen | | | 4.815 | 12 | –65.5 | –16 |
| R cerebellum declive | | | 4.478 | 30 | –59.5 | –22 |
| **L middle frontal gyrus (10)** | **153** | **<0.0001** | **4.600** | **–43.5** | **45.5** | **15.5** |
| L middle frontal gyrus | | | 3.929 | –36 | 39.5 | 11 |
| **R anterior cingulate gyrus** | **85** | **0.0029** | **4.365** | **1.5** | **12.5** | **39.5** |
| R anterior cingulate gyrus (32) | | | 4.122 | 9 | 15.5 | 33.5 |

Bold font indicates peak voxel.

**Supplementary Table 4. Brain Regions with Significant Responses to Imagining Odors > Smelling Clean Air**

| Left/Right Label (Brodmann Area) | Size (Voxels) | $p_{FWE}$ | t | MNI | | |
|---|---|---|---|---|---|---|
| | | | | x | y | z |
| **L insula anterior inferior cortex** | **3395** | **<0.0001** | **8.392** | **–40.5** | **5** | **–2.5** |
| L insula posterior short gyrus | | | 7.779 | –36 | –1 | 8 |
| L insula anterior short gyrus | | | 7.245 | –34.5 | 14 | 0.5 |
| **R insula anterior inferior cortex** | **3533** | **<0.0001** | **7.839** | **46.5** | **9.5** | **–4** |
| R precentral gyrus (44) | | | 7.744 | 45 | 0.5 | 6.5 |
| R insula middle short gyrus / inferior frontal gyrus (13) | | | 7.696 | 43.5 | 8 | 3.5 |
| **R putamen / piriform cortex** | **845** | **<0.0001** | **7.252** | **22.5** | **5** | **–5.5** |
| R pallidum | | | 5.762 | 12 | 3.5 | –7 |
| R putamen | | | 5.110 | 27 | –14.5 | 5 |
| **L precentral gyrus** | **193** | **<0.0001** | **6.637** | **–39** | **–16** | **39.5** |
| **L putamen / piriform cortex** | **598** | **<0.0001** | **5.995** | **–19.5** | **3.5** | **–7** |
| L pallidum | | | 4.836 | –12 | 6.5 | –7 |
| L putamen | | | 4.596 | –22.5 | 8 | 6.5 |
| **R posterior orbitofrontal gyrus** | **70** | **0.0283** | **5.116** | **22.5** | **20** | **–20.5** |
| R posterior orbitofrontal gyrus | | | 3.801 | 25.5 | 30.5 | –16 |

| | | | | | | |
|---|---|---|---|---|---|---|
| **R precentral gyrus** | **138** | **0.0003** | **5.056** | **40.5** | **–13** | **38** |
| **R middle frontal gyrus** | **180** | **<0.0001** | **4.960** | **39** | **44** | **6.5** |
| R middle frontal gyrus (10) | | | 4.429 | 45 | 47 | –1 |
| R middle frontal gyrus | | | 3.953 | 45 | 50 | 9.5 |
| **L cerebellum declive** | **137** | **0.0003** | **4.915** | **–19.5** | **–59.5** | **–25** |
| L cerebellum declive | | | 4.328 | –24 | –65.5 | –22 |
| L cerebellum / dentate nucleus | | | 4.318 | –16.5 | –59.5 | –32.5 |
| **R postcentral gyrus (3)** | **181** | **<0.0001** | **4.913** | **57** | **–16** | **27.5** |
| R postcentral gyrus (43) | | | 4.524 | 55.5 | –16 | 18.5 |
| R supramarginal gyrus (40) | | | 4.207 | 52.5 | –28 | 29 |
| **L anterior orbitofrontal gyrus** | **188** | **<0.0001** | **4.868** | **–24** | **38** | **–16** |
| L posterior orbitofrontal gyrus | | | 4.829 | –24 | 17 | –22 |
| L posterior orbitofrontal gyrus (11) | | | 4.729 | –19.5 | 27.5 | –23.5 |
| **R hippocampus / posterior temporal lobe** | **69** | **0.0305** | **4.624** | **25.5** | **–35.5** | **–4** |
| R parahippocampal gyrus / posterior temporal lobe | | | 3.824 | 18 | –38.5 | 3.5 |
| **L middle frontal gyrus (10)** | **407** | **<0.0001** | **4.581** | **–39** | **50** | **20** |
| L middle frontal gyrus | | | 4.577 | –37.5 | 45.5 | 27.5 |
| L middle frontal gyrus | | | 4.441 | –39 | 41 | 17 |
| **R anterior cingulate gyrus** | **97** | **0.0042** | **4.327** | **10.5** | **14** | **38** |
| R anterior cingulate gyrus (32) | | | 3.549 | 4.5 | 21.5 | 33.5 |
| **R supramarginal gyrus (40)** | **69** | **0.0305** | **4.189** | **57** | **–35.5** | **29** |

Bold font indicates peak voxel.

**Supplementary Table 5. Correlations Comparing the Right Piriform Decoding of Real and Imagined Odors Versus the Perceptual and Self-Report Measures of Odor Imagery Ability in the Full Sample (n = 44)**

| Variable | Right Piriform Decoding of Real Odors | | Right Piriform Decoding of Imagined Odors (Neural Measure of Odor Imagery Ability) | |
|---|---|---|---|---|
| | r | p | r | p |
| VOIQ | –0.196 | 0.2033 | 0.289 | 0.0571 |
| VFIQ | –0.079 | 0.6104 | 0.274 | 0.0722 |
| Interference Effect | –0.129 | 0.4025 | **0.352** | **0.0191*** |

VOIQ, Vividness of Olfactory Imagery Questionnaire[30]; VFIQ, Vividness of Food Imagery Questionnaire[20]. *$p < 0.05$.

**Supplementary Table 6. Correlations Comparing the Food Cue Reactivity Measures with Demographics, Adiposity, Food Liking, Hunger, and Consumption of Unhealthy Foods**

| Variable (Units) | Food Craving | | Food Intake | |
|---|---|---|---|---|
| | r | p | r | p |

| | | | | |
|---|---|---|---|---|
| Sex | 0.244 | 0.1062 | **0.365** | **0.0161*** |
| Age (yr) | –0.126 | 0.4101 | –0.098 | 0.5333 |
| Household Income | 0.143 | 0.3483 | –0.120 | 0.4423 |
| BMI (kg/m$^2$) | –0.270 | 0.0733 | 0.086 | 0.5824 |
| Body Fat (%) | –0.270 | 0.0726 | 0.131 | 0.4023 |
| Liking for Craving Stimuli | **0.521** | **0.0002**** | 0.180 | 0.2473 |
| Liking for Intake Stimuli | 0.174 | 0.2522 | **0.349** | **0.0217*** |
| Hunger | **0.497** | **0.0005**** | 0.282 | 0.0672 |
| DFS Score | 0.201 | 0.1859 | –0.076 | 0.6305 |
| Frequency of Consumption for Craving Stimuli | 0.187 | 0.2187 | 0.237 | 0.1253 |
| Frequency of Consumption for Intake Stimuli | 0.043 | 0.7775 | 0.078 | 0.6186 |

Males consumed more than females ($t_{41} = 2.511$). DFS, Dietary Fat and Free Sugar Short Questionnaire[118], American version. *p < 0.05; **p < 0.001.

**Supplementary Table 7. Correlations Comparing Adiposity Change Over One Year with Demographics, Olfactory Function, Food Liking, Consumption of Unhealthy Foods, and Physical Activity Change**

| Variable (Units) | Δ BMI | | Δ Body fat % | |
|---|---|---|---|---|
| | r | p | r | p |
| Sex | < 0.001 | 0.9980 | 0.094 | 0.5743 |
| Age (yr) | **–0.323** | **0.0348*** | –0.011 | 0.9428 |
| Household Income | 0.113 | 0.4720 | 0.116 | 0.4574 |
| Rose Odor Threshold | 0.013 | 0.9353 | 0.175 | 0.2627 |
| Cookie Odor Threshold | –0.015 | 0.9267 | –0.054 | 0.7309 |
| Liking for Craving Stimuli | 0.097 | 0.5366 | 0.152 | 0.3295 |
| Liking for Intake Stimuli | 0.273 | 0.0763 | 0.180 | 0.2480 |
| DFS Score | 0.045 | 0.7752 | –0.059 | 0.7079 |
| Frequency of Consumption for Craving Stimuli | 0.112 | 0.4765 | 0.146 | 0.3495 |
| Frequency of Consumption for Intake Stimuli | 0.119 | 0.4460 | 0.052 | 0.7412 |
| Change in IPAQ Weekly MET-minutes | –0.123 | 0.4316 | –0.238 | 0.1237 |

DFS, Dietary Fat and Free Sugar Short Questionnaire[118], American version; IPAQ, International Physical Activity Questionnaire[131]; MET, Metabolic Equivalents of Task. *p < 0.05; **p < 0.01.

**Supplementary Table 8. Participant Characteristics (N = 45)**

| Variable (Units) | Low BMI Group (n = 23) | | | High BMI Group (n = 22) | | | Group Difference | |
|---|---|---|---|---|---|---|---|---|
| | M | SD | Range | M | SD | Range | $t_{43}$ | p |
| Sex | 12 Male, 11 Female | | | 11 Male, 11 Female | | | N/A | N/A |
| Race | 9 White, 9 Asian, 2 Black/African American, 2 More than One Race, 1 Unknown or Preferred Not to Report | | | 12 White, 5 Black/African American, 3 Asian, 1 More than One Race, 1 Unknown or Preferred Not to Report | | | N/A | N/A |
| Ethnicity | 19 Not Hispanic/Latinx, 3 Hispanic/Latinx, 1 Unknown or Preferred Not to Report | | | 18 Not Hispanic/Latinx, 4 Hispanic/Latinx | | | N/A | N/A |
| Age (yr) | 25.6 | 5.8 | 19–37 | 28.4 | 6.0 | 18–42 | 1.588 | 0.1195 |
| Household Income | 4.2 | 1.8 | 1–8 | 3.7 | 1.9 | 1–8 | 0.972 | 0.3363 |
| Baseline Height (m) | 1.72 | 0.11 | 1.54–1.93 | 1.67 | 0.09 | 1.50–1.84 | 1.722 | 0.0922 |
| Baseline Weight (kg) | 64.48 | 11.62 | 45.90–85.00 | 86.31 | 23.51 | 66.00-158.00 | **3.975** | **0.0003*** |
| Baseline BMI ($kg/m^2$) | 21.62 | 1.67 | 18.32–24.28 | 30.82 | 7.00 | 25.25–53.44 | **6.123** | **< 0.0001**** |
| Baseline Body Fat % (Sex-Adjusted) | 0.81 | 0.15 | 0.50–1.20 | 1.33 | 0.30 | 0.92–2.17 | **7.627** | **< 0.0001**** |
| Change in Weight (kg) | 0.87 | 3.62 | –7.15–7.25 | 2.14 | 3.32 | –3.15–9.65 | 1.490 | 0.1438 |
| Change in BMI ($kg/m^2$) | 0.22 | 1.28 | –3.04–2.14 | 0.69 | 1.08 | –1.17–2.84 | 1.480 | 0.1465 |
| Change in Body Fat % (Sex-Adjusted) | 0.02 | 0.12 | –0.22-0.21 | 0.04 | 0.10 | –0.14–0.27 | 0.999 | 0.3238 |

Household income was dummy coded from 1–8 according to 2018 US Census Bureau income percentiles. Changes in adiposity reflect differences from the baseline to one-year follow-up sessions (n = 43). N/A, Not Applicable. *p < 0.001; **p < 0.0001.

**Supplementary Table 9. Test-Retest Reliability for the Questionnaires and Food Cue Reactivity Measures Completed at the Baseline and One-Year Follow-Up Sessions**

| Variable | ICC | 95% CI Lower | 95% CI Upper |
|---|---|---|---|
| VOIQ | 0.684 | 0.484 | 0.816 |
| VFIQ | 0.609 | 0.380 | 0.768 |
| VVIQ | 0.627 | 0.401 | 0.781 |
| DFS | 0.790 | 0.644 | 0.880 |
| Food Craving | 0.673 | 0.467 | 0.809 |
| Food Intake | 0.662 | 0.443 | 0.806 |

ICC, Intraclass correlation coefficient; VOIQ, Vividness of Olfactory Imagery Questionnaire[30]; VFIQ, Vividness of Food Imagery Questionnaire[20]; VVIQ, Vividness of Visual Imagery Questionnaire[17]; DFS, Dietary Fat and Free Sugar Short Questionnaire[118], American version.
